# Single-cell profiling of penta- and tetradactyl mouse limb buds identifies mesenchymal progenitors controlling digit numbers and identities

**DOI:** 10.1101/2024.06.03.597099

**Authors:** Victorio Palacio, Anna Pancho, Angela Morabito, Jonas Malkmus, Zhisong He, Geoffrey Soussi, Rolf Zeller, Barbara Treutlein, Aimée Zuniga

## Abstract

The cellular mechanisms controlling digit numbers and identities have remained elusive. Profiling of wild-type (pentadactyl) and *Grem1* tetradactyl mouse limb buds identifies cellular changes affecting two limb bud mesenchymal progenitor (LMP) populations. In mutant limb buds, the anteriorly biased distribution of peripheral LMPs (pLMPs) is lost and the population expanded, while the distal-posterior LMP (dLMP) population is reduced from early stages onward. Analysis of LMP signature genes in wildtype and mutant mouse limb buds with digit loss or gain establishes that pLMPs are positively regulated by BMP signaling, while dLMPs require GREM1-mediated BMP antagonism. dLMPs encompass digit progenitors and altering their population size prefigures changes in digit numbers. The anteriorly biased pLMP distribution is linked to digit asymmetry as loss of this bias in tetradactyl mouse and pig limb buds underlies middle digit symmetry and identity loss. This study indicates that variable spatial *Grem1* expression in mutant and evolutionary diversified limb buds tunes BMP activity, impacting both LMP populations in a complementary manner.

## Introduction

Elucidating the molecular events underlying vertebrate limb buds is shedding light both on our understanding of human limb congenital malformation and evolutionary limb diversification. One of the most important signaling molecules for limb development is the Sonic Hedgehog (SHH) morphogen. SHH is expressed by the posterior mesenchymal organizer and required for both antero-posterior (AP) patterning of limb skeletal elements, and survival and proliferative expansion of limb bud mesenchymal cells.^1-3^ Its temporally controlled genetic inactivation in mouse limb buds reveals that *Shh* acts early and short range to directly specify posterior digits during onset of limb bud development (E9.75-E10.0).^1,3^ Previous studies indicated that posterior digit specification depends on the time mesenchymal cells are exposed to *Shh*, while the anterior digit 2 is specified by long-range SHH signaling.^4-6^ BMP signaling is also required for digit development^7-11^ and one key event is the activation and upregulation of the BMP antagonist *Gremlin1* (*Grem1*) by BMPs.^12^ Concurrently, SHH upregulates *Grem1* in the posterior-distal limb bud mesenchyme^13,14^, which results in establishment of the self-regulatory SHH/GREM1/AER-FGF feedback signaling system that promotes cell survival and proliferation in concert with WNT signaling.^1,12,14-18^ An integral part of this self-regulatory signaling system is the spatially dynamic transcriptional regulation of *Grem1*, which preferentially antagonizes SMAD4-mediated BMP signal transduction in the posterior and distal limb bud mesenchyme.^19,20^ In *Grem1*-deficient mouse limb buds, the SHH/GREM1/AER-FGF feedback signaling system is disrupted, which cause mesenchymal apoptosis and formation of only 3 very rudimentary digits.^21,22^ The spatial robustness of *Grem1 cis*- regulation is paramount to normal limb and digit development as progressive genetic reduction of its spatial domain and posterior expression bias causes transition from wildtype pentadactyly (5 digits) to stable tetradactyly (4 digits) via intermediate and variable anterior digits fusions.^23^ In *Grem1* tetradactyl limbs, the loss of an anterior digit is paralleled by a shift of the median axis from digit 3 in wildtypes (mesaxonic) to a paraxonic position in-between the symmetrical middle digits (dotted lines, Figure 1A). The paraxonic limb axis is the defining feature underlying establishment of the unguligrade posture during evolutionary diversification of Artiodactyl limbs.^24,25^ During handplate (autopod) development, the AP positions and alternating sequence of digit and interdigit tissues are determined by a self-regulatory and robust three-node BMP/SOX9/WNT Turing-system.^26,27^ Digit identities are determined late by signals from the interdigit and AER acting on the phalanx forming region (PFR).^28-30^ The activity of the PFR regulates the number and length of phalanges formed, which is the main morphological feature of digit identities in addition to their AP positions.^29,31^ Currently there is a significant knowledge gap in understanding how the early and direct specification of posterior digits by SHH^3^ is linked to establishment of the definitive digit pattern and identities during autopod development. This early specification must trigger molecular changes in target limb bud mesenchymal progenitor (LMP) cells that are propagated during progression of limb bud outgrowth and autopod development. Indeed, there is molecular evidence for intrinsic cellular memories as the length of Shh exposure specifies the identities of the *Shh* descendants that contribute to the posterior-most digits d4 and d5.^6^ Similarly, transient exposure of chicken wing bud mesenchyme to AER-FGF signaling triggers an intrinsic gene regulatory network (GRN) that controls the proliferative expansion of the distal progenitors giving rise to the autopod.^32^ Cell lineage analysis in mice shows that the *Hoxa13* transcriptional regulator is the earliest marker for LMPs that give rise to all skeletal elements of the autopod with exception of the proximal-most carpal elements.^33^ Recently, Markman and co-authors^34^ have combined single cell profiling of mouse limb buds with lineage analysis of *Msx1*-positive LMPs to study the complex spatio-temporal dynamics of skeletal morphogenesis. They show that the *Msx1*^+^ progenitors progressively differentiate into the *Sox9*-positive osteochondrogenic progenitors (OCPs) that contribute to all skeletal primordia. Both mouse and human limb bud single cell studies show that *Msx1*^+^ progenitors are highly heterogeneous and can be subdivided into populations with distinct locations based on additional markers.^34,35^ For example, one *Msx1^+^* population contributes to distal skeletal progenitors based on a set of genes expressed in the autopod that include *5’Hox* genes, *Lhx2/9*, *Msx1/2*, *Rdh10*, *Sp9*, and *Tfap2a/b*.^34,35^ An important feature of the spatio-temporal kinetics governing this process is the progressive yet at the same time synchronous transition of *Msx1*^+^ progenitors to both proximal and distal skeletal progenitor populations. How this heterogenous population gives rise to a given number of digits with specific identities remains unknown. To bridge this knowledge gap and gain functional insights, we combined comparative high resolution gene expression analysis of wildtype and tetradactyl mouse limb buds at early stages with lineage tracing. This identifies two novel LMP populations present from early limb bud outgrowth onward, that are spatially complementary and appear to full-fill distinct functions in specification of digit numbers and identities. Their differential regulation by GREM1-mediated local modulation of BMP activity impacts the size and spatial distribution of these populations in an opposing manner. *Grem1*-dependent alterations of these LMP populations underlie changes in digit numbers and identities in limb congenital malformations and evolutionary digit reductions.

**Figure 1.**
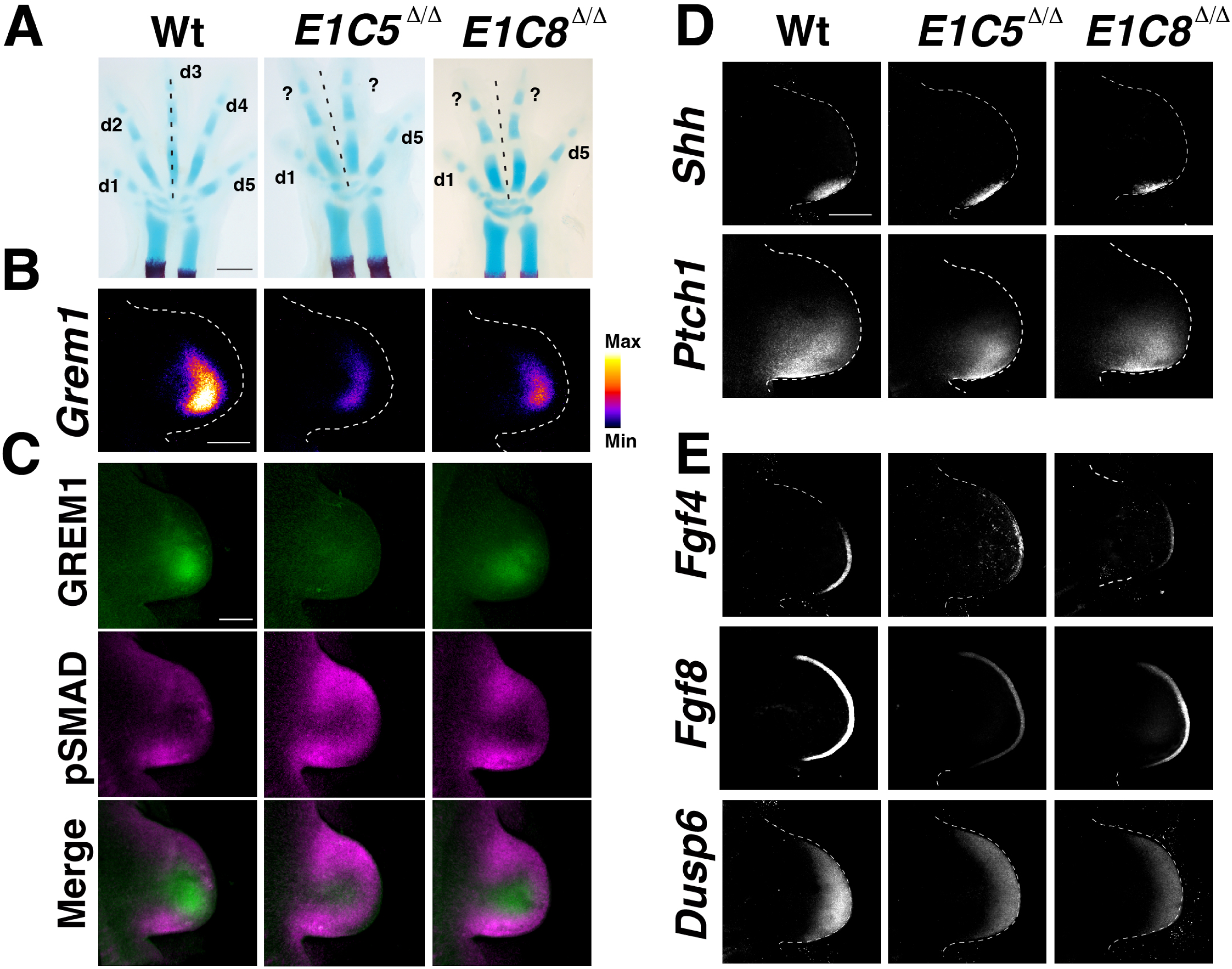
Analysis of the feedback signaling system in wildtype and tetradactyl mouse forelimb buds. (A) Skeletal analysis of wildtype (Wt) pentadactyl, *E1C5* ^Δ/Δ^ and *E1C8* ^Δ/Δ^ tetradactyl forelimbs at E14.5. Cartilage is stained blue, the ossification centers in radius and ulna in red. Digits are indicated from anterior (d1) to posterior (d5). Digits with uncertain identities are labelled by “?”. The dotted line indicates the median axis in both pentadactyl and tetradactyl limbs. Per genotype, minimally n=3 embryos independent biological replicates were analyzed. Scale bar: 1 mm. (B) Analysis of the spatial distribution of *Grem1* transcripts by whole mount RNA-FISH in forelimb buds of all three genotypes at E10.75 (37-39 somites, n≥3 replicates per genotype). The transcript levels are shown as intensity values using the fire look up table of FIJI. (C) Merge of GREM1 and pSMAD activity by whole mount immunofluorescence in forelimb buds of all three genotypes at E10.75 (n≥3 replicates per genotype). Scale bar: 200 μm. (D) Analysis of *Shh* and *Ptch1* expression by whole mount RNA-FISH in forelimb buds at E10.75 (n≥3 replicates per genotype). (E) Analysis of AER-FGF (*Fgf4* and *Fgf8*) expression and mesenchymal FGF signal transduction sensed by *Dusp6* expression in forelimb buds at E10.75 (n≥3 replicates per genotype). All forelimb buds are oriented with anterior to the top and posterior to the bottom. Scale bar: 300μm. Relates to Figure S1.

## Results

### Comparative analysis of wild-type and *Grem1* tetradactyl mouse forelimb buds identifies the lineage alterations underlying the anterior digit loss

Two different *Grem1* alleles lacking 4 of 7 limb bud CRM enhancers, namely *EC1CRM5*^Δ/Δ^ (*E1C5* ^Δ/Δ^)^23^ and *EC1CRM8* ^Δ/Δ^ (*E1C8*^Δ/Δ^) result in tetradactyl limb phenotypes with loss of an anterior digit (Figure 1A). Analysis of *E1C5*^Δ/Δ^ and *E1C8*^Δ/Δ^ mouse forelimb buds at embryonic day E10.75 (37-39 somites) by fluorescent whole mount HCR^TM^ RNA *in situ* hybridization (RNA-FISH) and immunofluorescence^36^ shows that the tetradactyl phenotype is caused by a remarkable spatial reduction of the *Grem1* transcript and protein expression domains (Figures 1B, 1C). In particular, the posterior bias and distal-anterior expansion of *Grem1* expression is disrupted in tetradactyl limb buds (Figure 1B).^23^ Spatial reduction in GREM1 results in complementary expansion of pSMAD1/5/9 (pSMAD) activities that points to increased BMP signal transduction (Figures 1C and S1A).^37,38^ Furthermore, *Shh* expression and the upregulation of *Ptch1* in SHH responsive cells is reduced (Figure 1D). Moreover, AER-*Fgf* expression and FGF signal transduction in the distal-most mesenchyme are also reduced as indicated by the FGF target *Dusp6*^39^ and lowered pERK levels^40^ (Figure 1E and S1B). This establishes that the self-regulatory SHH/GREM1/AER-FGF feedback signaling system^12,14-16^ remains intact but is scaled down in both *E1C5*^Δ/Δ^ and *E1C8*^Δ/Δ^ limb buds due to the spatial restriction and reduction of GREM1-mediated BMP antagonism. These well-defined tetradactyl phenotypes provide a unique opportunity to study the molecular and cellular alterations that underlies tetradactyly and explore the potential relevance for evolutionary digit reductions.

To identify the anterior digit lost in *Grem1* tetradactyl limbs, genetic lineage analysis was performed in wildtype and *E1C5^Δ/Δ^* embryos using specific CRE drivers to permanently activate a fluorescent cell lineage marker in either the posterior or anterior forelimb bud mesenchyme (Figure 2; single channels of the same limb buds are shown in Figures S2A, S2B). Posterior cell lineages were traced using the knock-in *Shh*^GFP*Cre*^ allele in combination with conditional activation of the ROSA26^LSL*-tdTomato*^ reporter in *Shh*-descendant cells (Figure 2A, 2B).^6,41^ In wild-type forelimb buds, the *Shh* lineage contributes to the posterior half of the autopod, with most descendants in digits d4 and d5 and lesser contributions to interdigit 3 and the posterior part of digit d3 (arrowhead and insets, Figure 2A).^6^ In *E1C5*^Δ/Δ^ forelimb buds, the posterior lineage contribution is comparable to wildtype with highest contributions to digits d5 and d4 and fewer cells to the interdigit between digit d4 and the 3^rd^ digit from posterior (d3*, Figure 2B; compare insets between Figures 2A, 2B). Complementarily, the anterior lineage was traced using the tamoxifen-inducible *Alx4*-*Cre*^ERT2^ transgene that is expressed in the anterior peripheral mesenchyme up to the limb apex^42^ in combination with the conditional ROSA26^LSL-*EGFP*^ reporter.^43^ The anterior lineage was labelled by activating the *Alx4*-*Cre*^ERT2^ transgene at ∼E10.5-10.75 (Figure 2C, 2D). In wild-type forelimb buds, the *Alx4*-*Cre*^ERT2^ lineage contributes to the anterior half of the autopod with descendants detected up to the anterior part of digit d3 (arrowhead and insets, Figure 2C).^43^ In *E1C5^Δ Δ^* forelimb buds, the lineage*^/^*is more restricted due to an anterior shift of the lineage boundary (arrowhead, Figure 2D) and descendants are detected in the anterior-most digit d1, interdigit 1 and the 3^rd^ digit from posterior (d3*; insets, Figure 2D). Both lineages of two age- and shape-matched forelimb buds from different embryos were virtually overlapped to visualize the AP boundary (Figure 2E). Whereas in wildtype forelimb buds, both lineages contribute to digit 3 (upper panels, Figure 2E), in *Grem1* tetradactyl limb buds, both lineages contribute to the third digit from the posterior, identifying this digit as d3 while digit d2 is lost (lower panels, Figure 2E). This is corroborated by the detection of occasional *Sox9*-positive condensations in the interdigit space between digit d1 and d3 in *Grem1* tetradactyl limb buds (Figure S2E). However, these ectopic condensations are transient as they do not contribute to the digit skeleton (Figure 1A). In addition, this analysis establishes that in *Grem1* tetradactyl mouse limbs the asymmetry of middle digits d3 and d4 is lost, which contrasts with wild-type pentadactyly.

**Figure 2.**
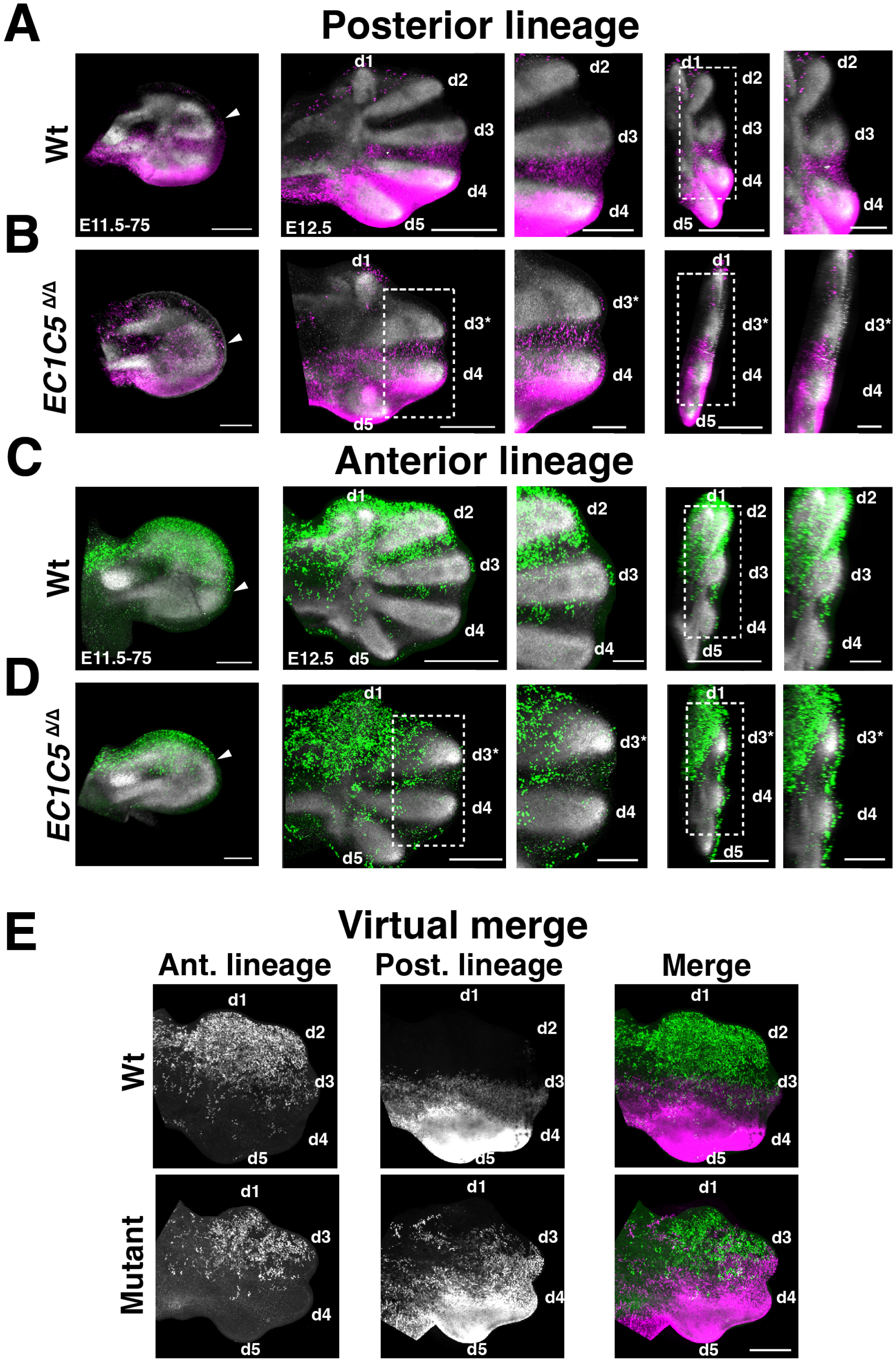
Posterior and anterior lineage analysis of pentadactyl and *Grem1* tetradactyl forelimb buds. (A) Tracing the posterior lineage by *Shh*^CRE^-mediated permanent activation of the ROSA26tdTomato reporter (magenta). Representative forelimb buds are shown. Left panel shows a wildtype forelimb bud at E11.5-E11.75 (n=6/6). Arrowhead indicates the anterior boundary of the posterior lineage. Middle and right panels show wildtype forelimb buds at E12.5 (n=6/6). Enlargements show optical sections to better reveal the posterior lineage between digits d2 to d4. Scale bar (left panel): 200 µm. (B) Tracing the posterior lineage in tetradactyl *E1C5* ^Δ/Δ^ forelimb buds. Left panel: *E1C5* ^Δ/Δ^ forelimb bud at E11.5-E11.75. Arrowhead indicates the anterior boundary of the posterior lineage, which is shifted in comparison to the wildtype (n=3/4). Middle and right panels: *E1C5* ^Δ/Δ^ forelimbs at E12.5. Enlargements show optical sections to better reveal the posterior lineage between digits d3*and d4. The lineage extends to the middle of d3* (n=5/6). (C) Tracing the anterior lineage by tamoxifen-mediated induction of the *Alx4*-Cre^ERT2^ transgene around E10.5, which activates the Rosa26eGFP reporter (green). Representative forelimb buds are shown. Left panel shows a wildtype forelimb bud at E11.5-E11.75 (n=6/6). Arrowhead indicates the posterior boundary of the anterior lineage. Scale bar (left panel): 200 µm. Middle and right panels show a wildtype forelimb at E12.5 (n=5/6). Enlargements show optical sections to better reveal the anterior lineage. (D) Anterior lineage analysis of tetradactyl *E1C5* ^Δ/Δ^ forelimb buds. Left panel shows a *E1C5* ^Δ/Δ^ forelimb bud at E11.5-E11.75. Arrowhead indicates the posterior boundary of the anterior lineage. Middle and right panels show a *E1C5* ^Δ/Δ^ forelimb at E12.5-75. Enlargements show optical sections to better reveal the region between digit d3* and d4. Scale bars for all middle and right panels: 500 µm (overviews), 250 µm (insets). (E) Virtual merge of the posterior and anterior lineages from two different limb buds at E12.5-75. Left and middle panels show single gray channels showing the anterior and posterior lineages after transformation. Right panels show the virtual merge of the anterior and posterior lineages. Scale bar: 300 µm. All limb buds are oriented with anterior to the top and posterior to the bottom, proximal to the left and distal to the right. Relates to Figure S2.

### Single cell RNA-sequencing identifies two early LMP populations altered in an opposing manner in the distal mesenchyme of *Grem1* tetradactyl forelimb buds

Comparative scRNA-seq analysis of *Grem1* tetradactyl and wild-type forelimb buds at E10.75 (37-39 somites) should reveal the early molecular and cellular changes that are linked to loss of digit 2 and middle digit asymmetry. This early stage was chosen as no morphological differences between wild-type and tetradactyl limb buds are apparent prior to autopod development (≥E11.25), and the LMPs contributing to autopod and digit development are specified during this developmental period.^34,44^ Furthermore, the progression of limb bud outgrowth and patterning is predominantly controlled by the SHH/GREM1/AER-FGF feedback signaling system at this stage (Figure 1 and S1).^12,14-16,45^ Comprehensive single cell datasets from wildtype, *E1C5* ^Δ/Δ^ and *E1C8* ^Δ/Δ^ forelimb buds were generated, verified and after initial processing, 38460 mesenchymal cells from the three genotypes were pooled (Wt: 12164 cells, *E1C5* ^Δ/Δ^: 11946 cells, *E1C8* ^Δ/Δ^: 14350 cells; see Methods and Figure 3A). Unsupervised analysis identified 12 distinct cell clusters (Figure 3B), which are composed of cells from all three genotypes (Figure S3A). The heatmap with the top 10 highest expressed genes for each cluster contains a high number of genes with known functions during limb bud development and/or spatially restricted expression patterns (Figure S3B).

**Figure 3.**
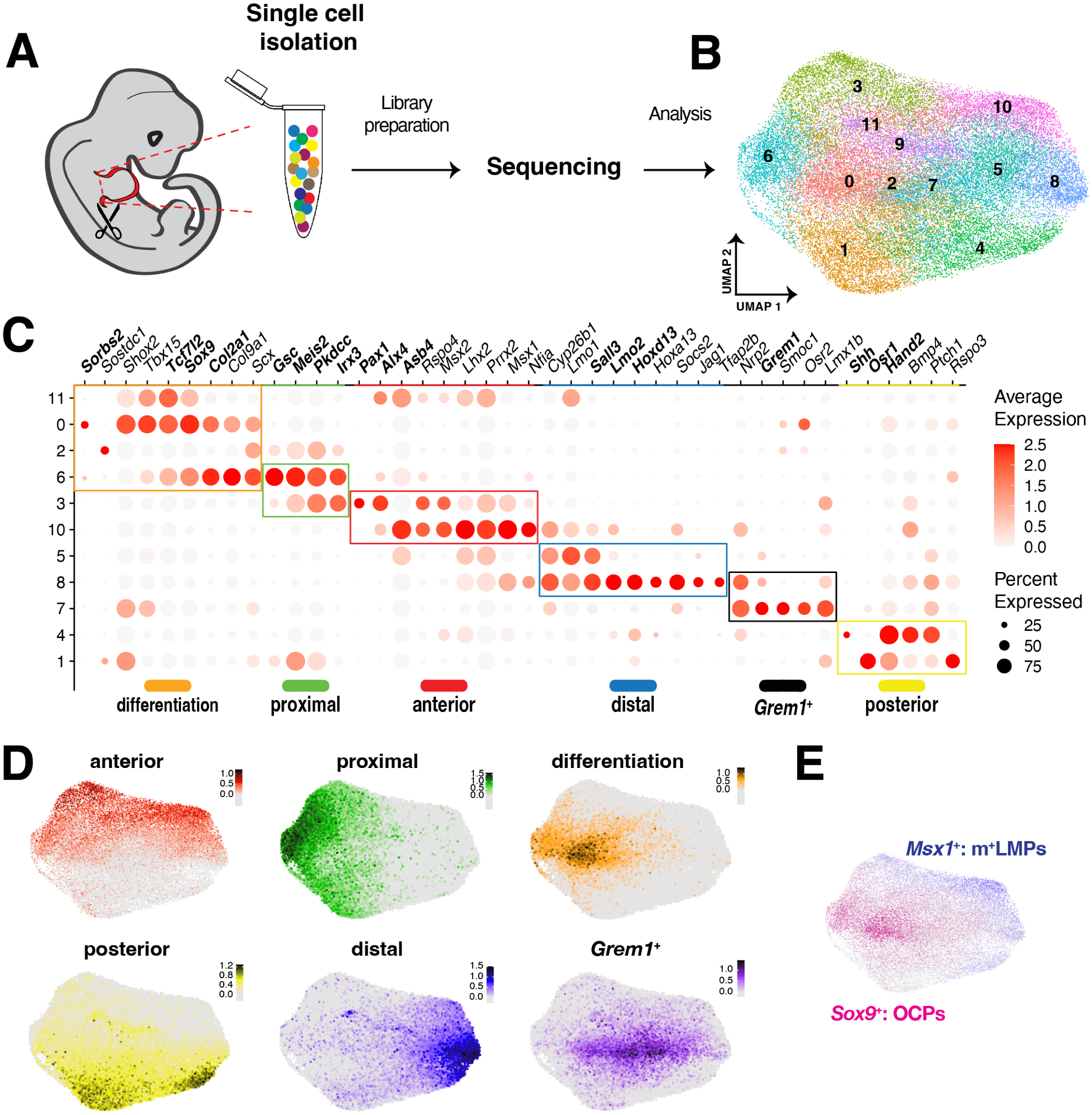
Comparative single cell RNA sequencing of wildtype pentadactyl and *Grem1* tetradactyl forelimb buds (E10.75). (A) Scheme showing the experimental procedure. Forelimb bud pairs from wildtype, *E1C5* ^Δ/Δ^ and *E1C8* ^Δ/Δ^ embryos (n=3 biological replicates per genotype) were dissected and dissociated into single cells. After quality check, libraries were prepared and sequenced. (B) UMAP embedding of the pooled scRNA-seq data from all three genotypes. (C) Dot plot showing the expression of different marker genes for the 12 UMAP cell clusters. Genes were ordered according to known patterns of expression in mouse limb bud at E10.75. Colored boxes and lines under gene names indicate the areas in limb buds where the marker genes are prominently expressed. (D) UMAP embedding of scores using positional markers. Genes used in respective scores were: anterior score: *Alx4*, *Asb4*, *Pax1*; posterior score: *Hand2*, *Osr1*, *Shh*; proximal score: *Gsc, Irx3, Meis2, Pkdcc*; distal score: *Hoxd13*, *Sall3*, *Lmo2*; differentiation score: *Col2a1*, *Sorbs2*, *Sox9*, *Tcf7l2*; *Grem1*^+^ score: *Grem1* expressing cells. These genes are indicated in bold in panel C. (E) UMAP embedding of *Msx1* to show the distribution of m^+^LMPs, and of *Sox9* for the distribution of OCPs. Relates to Figure S3.

To associate clusters to specific spatial regions in the limb bud mesenchyme we selected some of these genes to generate a dot plot (Figure 3C). This dot plot representation shows the differential expression of marker genes between clusters, which is either due to differences in transcripts levels or fraction of expressing cells (Figure 3C). This allowed the creation of a score for the relevant spatial regions in limb buds using specific genes as spatial indicators (Figure S3C). Visualization of these scores by UMAP projections shows that unsupervised clustering accurately reproduces the spatial organization of the limb bud mesenchyme (Figure 3D). For example, cell clusters 8 and 5 are located distally, while cell clusters 4 and 10 are located in the posterior-distal and anterior-distal mesenchyme respectively. These distal clusters are of particular interest as they are part of the peripheral *Msx1*^+^ progenitor population (m^+^LMPs, Figure 3E), which also encompasses the progenitors of the future autopod.^34^

Comparison of the population sizes for these distal clusters (Figure 4A) reveals that for cluster 4 and 10, the percentages of cell numbers are proportionally increased in both mutant limb buds. In contrast, cell numbers are reduced in the distal-most cluster 8 in both *E1C5* ^Δ/Δ^ and *E1C8* ^Δ/Δ^ limb buds in comparison to wildtype limb buds (Figure 4A). The reduced cluster 8 LMP population in *Grem1* tetradactyl limb buds uncovers a potential cellular link to the subsequent loss of digit d2 (Figure 2). Volcano plot visualization of differentially expressed genes (DEGs) for cluster 8 identified both down- (blue) and upregulated (red) DEGs in *Grem1* tetradactyl forelimb buds (E10.75, Figure 4B and S4A). In cluster 8, *Msx1* is the most upregulated DEG, which is corroborated by UMAP projection and bar plot analysis, which shows that the m ^+^LMP population is increased predominantly in the peripheral and distal clusters of *Grem1* tetradactyl limb buds (Figure 4C; left panels Figure S4E). In addition, several of the other DEGs upregulated in mutant limb buds are expressed in the peripheral mesenchyme and are anteriorly biased in wildtype limb buds. These DEGs include *Id1*, *Id2* and *Msx2* that are established sensors of changes in BMP signaling^46-48^, while *Lhx2*, *Asb4^49^*, *Prrx1*/*2*^50^, and *Rspo4*^51^ are direct targets of SMAD4-mediated BMP signal transduction (Figure 4B and S4A).^20^ Among these, *Lhx2*, *Asb4*, *Msx2* and *Rspo4* were designated as signature genes for a peripheral LMP population (pLMPs) that is enlarged in *Grem1* tetradactyl forelimb buds (Figures 4D and S4A, S4C). UMAP projections and bar plots show that the increase in pLMPs occurs in distal and distal-posterior clusters of mutant limb buds (arrowheads Figure 4D; middle panels Figure S4E). In contrast, the expression of DEGs downregulated in cluster 8 is mostly restricted to distal clusters in *Grem1* tetradactyl forelimb buds (Figure 4A, S4A and S4D). Therefore, seven genes expressed predominantly in cluster 8, including *Tfap2b*, *Jag1* and *Hoxa13*, were designated as signature genes for this distal-posterior LMP population (dLMPs) that is reduced in *Grem1* tetradactyl forelimb buds (Figures 4E, S4D and right panels Figure S4E). The dLMP population is reduced more in *E1C5*^Δ/Δ^ than *E1C8*^Δ/Δ^ limb buds (arrowheads Figure 4E), which correlates with the difference in downscaling the SHH/GREM1/AER-FGF feedback signaling system in the two types of tetradactyl limb buds (Figure 1 and S1). In summary, this unbiased comparative scRNA-seq analysis identifies pLMPs and dLMPs as two LMP populations with distinct molecular signatures that are altered in an opposing manner in response to reduced GREM1-mediated BMP antagonism. All pLMPs and dLMPs express *Msx1*, which accounts for the majority of m^+^LMPs in forelimb buds at E10.75.

**Figure 4.**
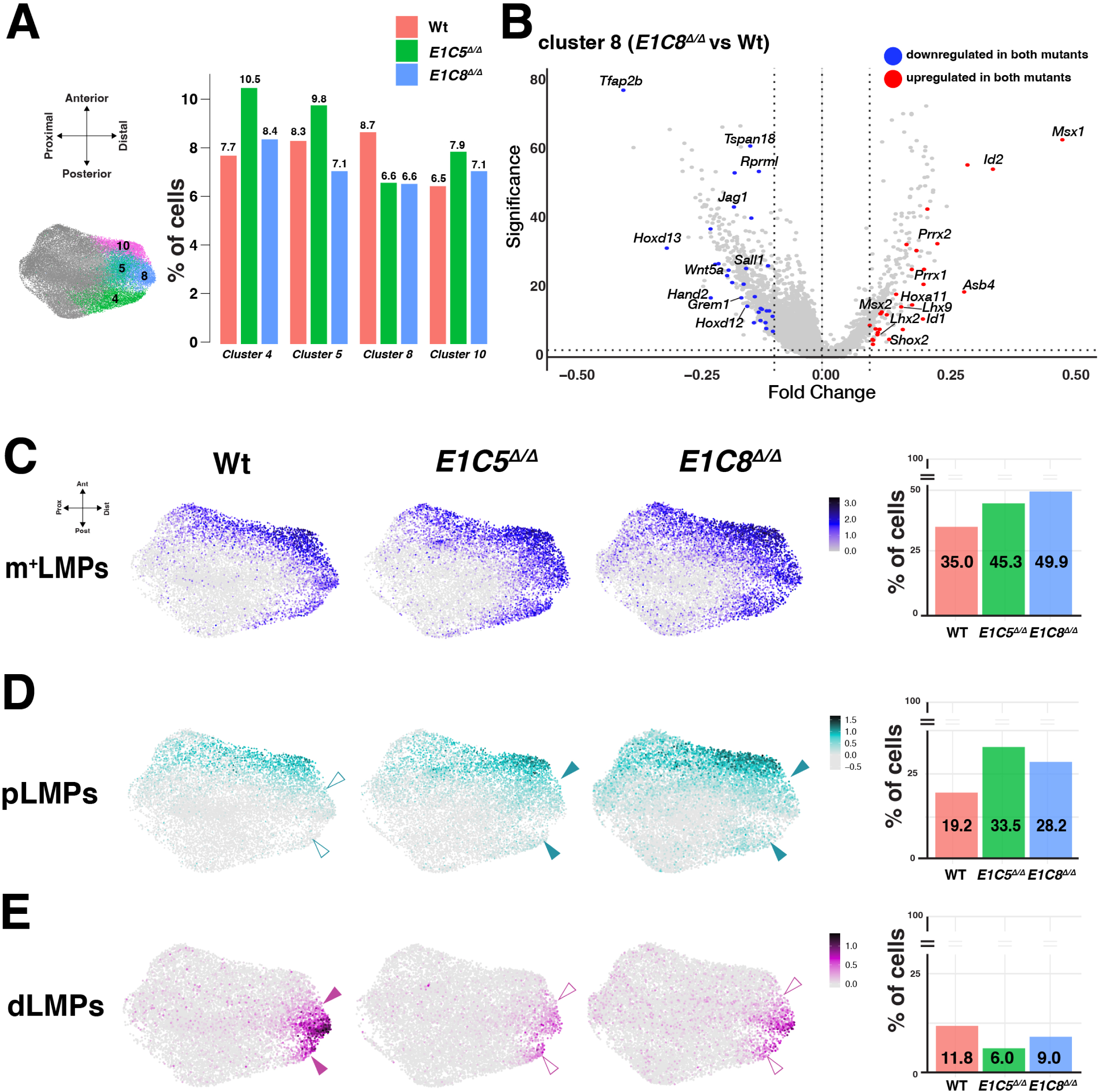
Identification of distinct LMP populations altered in both *Grem1* tetradactyl limb buds (E10.75). (A) UMAP projections show the distal clusters used for generating bar plots. Bars representing each of the four distal clusters show the percentage of cells in relation to all mesenchymal cell for each genotype. Cluster 4: posterior-distal; cluster 5 distal; cluster 8: distal-posterior, cluster 10: anterior-distal. (B) The volcano plot shows the differentially expressed genes (DEGs) in cluster 8 by comparing *E1C8* ^Δ/Δ^ to wildtype limb buds. DEGs indicated in blue are downregulated in both *E1C5* ^Δ/Δ^ (Figure S4) and *E1C8* ^Δ/Δ^ limb buds. Conversely, DEGs shown in red are upregulated in both *Grem1* tetradactyl limb buds. Fold changes (x-axis) are shown as Log2 values. Significance (y-axis) is Log2 of the adjusted p-values. (C-E) UMAP embedding scores for the three LMP populations in wildtype, *E1C5* ^Δ/Δ^ and *E1C8* ^Δ/Δ^ limb buds. Signatures for m^+^LMPs: *Msx1*; pLMPs: *Asb4*, *Lhx2*, *Msx2*, *Rspo4* (Figure S4B); dLMPs: *Tfap2b*, *Jag1*, *Hoxa13*, *Gja3*, *Pdzd2*, *Tspan18*, *Rprml* (Figure S4C). White arrowheads show downregulation while colored arrowheads show upregulation. Right-most panels: bar plots show the percentage of the different LMP cells in relation to all cells in wildtype, *E1C5* ^Δ/Δ^ and *E1C8* ^Δ/Δ^ limb buds. Relates to Figure S4.

### The reduction of dLMPs and concurrent increase of pLMPs points to a problem in correct specification of both population sizes in *Grem1* tetradactyl limb buds

To gain insight into the spatial alterations of LMP populations, key signature markers for each population were analyzed by RNA-FISH (Figures 5, S5). While *Msx1* expression is anteriorly biased in wildtype limb buds, its domain broadens in the distal-posterior mesenchyme in *E1C5* ^Δ/Δ^ and *E1C8* ^Δ/Δ^ limb buds (left panels Figure 5A, S5A). This expansion corresponds with the increased size of the m^+^LMP population (Figure 4C). Nevertheless, the *Msx1* and *Sox9* expression domains, which mark OCPs and the forming skeletal primordia,^34^ remain largely complementary in all three genotypes (right panels, Figure 5A, S5A). Two transcriptional regulators, *Lhx2^52^* and *Asb4*^53^ (Figure S4B), were chosen as signature genes to assess the spatial distribution of the pLMP population (Figures 4D, 5B). In contrast to the anterior bias in wildtype limb buds, *Lhx2* is upregulated and distal-posteriorly expanded in *E1C5* ^Δ/Δ^ and *E1C8* ^Δ/Δ^ forelimb buds (left panels, Figures 5B, S5B). Furthermore, the domain of *Asb4*^+^ pLMPs is enlarged (*E1C5* ^Δ/Δ^) or shifted distally (*E1C8* ^Δ/Δ^) respectively in *Grem1* tetradactyl limb buds (right panels Figure 5B, S5B). For dLMPs, the Notch ligand *Jag1* and the *Hoxa13* and *Tfap2b* transcriptional regulators were chosen as their expression is restricted to the posterior-distal mesenchyme (Figures 5C, 5E, S5C). In *E1C5* ^Δ/Δ^ and *E1C8* ^Δ/Δ^ limb buds, the spatial expression domains of both *Jag1* and *Tfap2b* are reduced, which matches the reduction of the dLMP population by UMAP projection (Figure 5C, S5C). Overlapping the expression domains of *Jag1* and *Lhx2,* and *Tfap2b* and *Asb4* respectively in the same limb bud reveals that the reduction of the dLMP population is accompanied by a complementary distal-posterior increase of the pLMP population in *Grem1* tetradactyl limb buds at E10.75 (Figure 5D).

**Figure 5.**
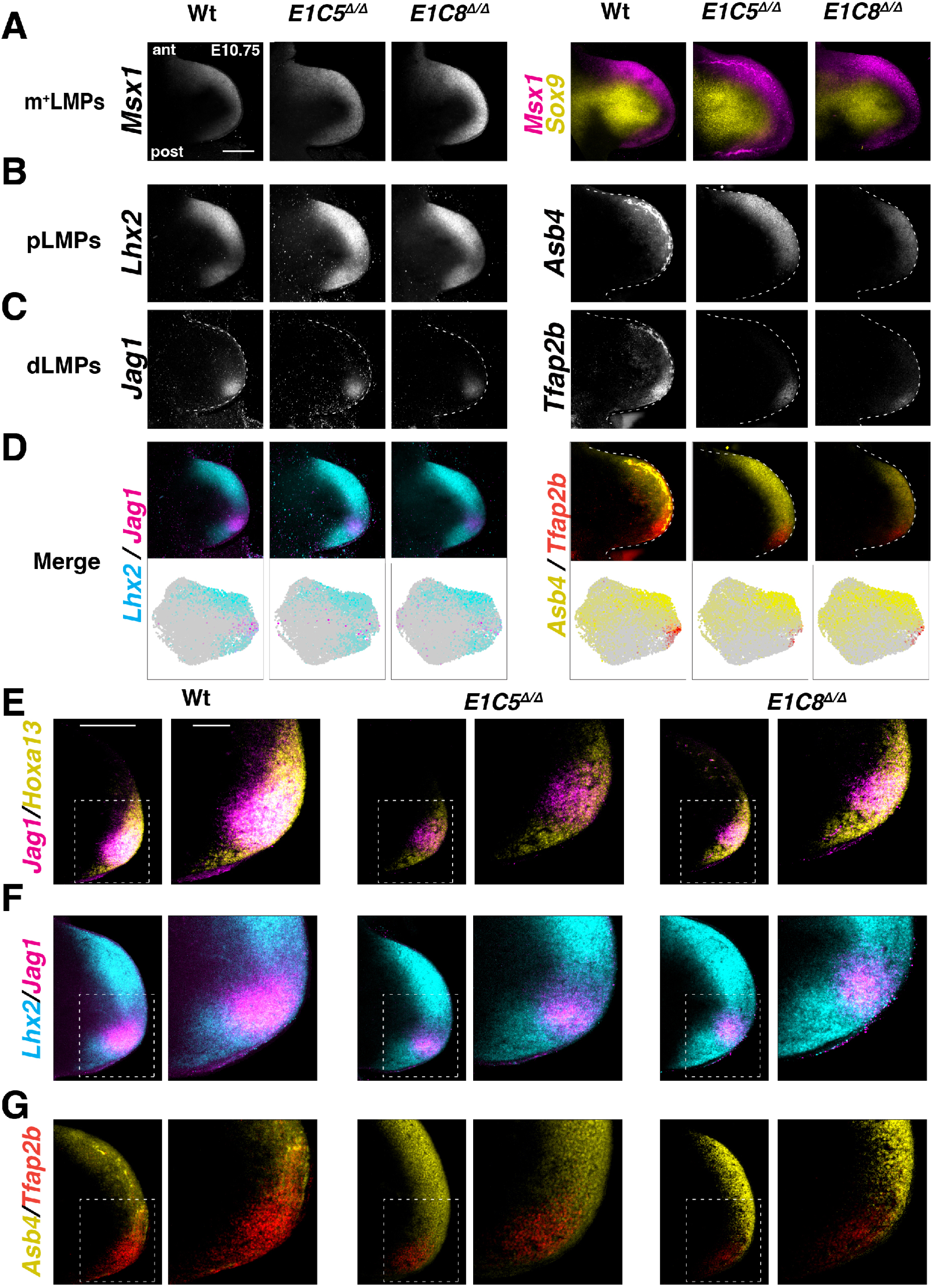
Opposing alterations and mixing of pLMPs with dLMPs point to a problem in specifying the dLMP population size in *Grem1* tetradactyl limb buds. Whole mount RNA-FISH analysis of wildtype and *Grem1* tetradactyl forelimb buds. All forelimb buds (E10.75) and enlargements are oriented with anterior to the top and posterior to the bottom. (A) Spatial distribution of m^+^LMPs (*Msx1*) and OCPs (*Sox9*) in wildtype and *E1C5* ^Δ/Δ^ and *E1C8* ^Δ/Δ^ limb buds. (B-D) Spatial distribution of pLMPs (*Lhx2* and *Asb4*) and dLMPs (*Jag1* and *Tfap2b*) in wildtype, *E1C5* ^Δ/Δ^ and *E1C8* ^Δ/Δ^ limb buds. (D) Upper panels left: merge of *Lhx2* (cyan) with *Jag1* (magenta). Right: merge of *Asb4* (yellow) with *Tfap2b* (red). Lower panels left: UMAP embeddings of *Lhx2* and *Jag1* (left). Right: *Asb4* and *Tfap2b*. (E-G) Medial optical sections (6.9μm) through the center of the dLMP domain shows the spatial expression at close to cellular resolution. (E) Overlap showing the *Jag1* domain nested within the *Hoxa13* domain. (F) Merge of *Lhx2* and *Jag1*-expressing LMPs. (G) Overlap of *Asb4* and *Tfap2b*-expressing LMPs. Scale bars: 300μm. Relates to Figure S5.

Optical sections of wildtype limb buds at close to cellular resolution show that the *Jag1* domain is nested within the larger *Hoxa13* domain (Figure 5E). While nesting is maintained in *Grem1* tetradactyl limb buds, both domains appear less defined and patchy (middle and right panels, Fig. 5E). The merge of *Jag1* with *Lhx2* and *Tfap2b* with *Asb4* in wildtype limb buds reveals the well-defined domains with some overlap of pLMPs and dLMP in the boundary regions of their respective domains (Figure 5F, 5G). In *Grem1* tetradactyl limb buds, *Lhx2*^+^pLMPs are mixed with the reduced *Jag1*^+^dLMP population in a salt and pepper pattern (Figure 5F). This intermingling of LMPs and reduction of dLMPs is also apparent from overlapping *Tfap2b*^+^dLMPs with the posteriorly expanded *Asb4*^+^pLMP population (Figure 5G). In summary, the RNA-FISH analysis complements the single cell analysis and shows that dLMPs and pLMP distributions are spatially distinct and largely complementary in wildtype limb buds at E10.75 (Figure 5D, F, G). By contrast, in *Grem1* tetradactyl forelimb buds, the dLMP population is reduced and less distinct and forms heterogeneous domain, in which dLMPs are mixed with pLMPs. The GREM1-dependent specification of dLMPs coincides with the prominent gap separating the larger anterior from the smaller posterior pLMP domain in wildtype forelimb buds at E10.75 (Figures 5B, 5D, 5F). Analysis of forelimb buds between E10.5 (35-36 somites) and E11.25 (43-44 somites; Figure 6) shows that dLMPs are already detected in forelimb buds at E10.5 (Figure 6A, 6B). Their spatial domain is reduced from the earliest stage onward and persists in *Grem1* tetradactyl forelimb buds (E11.5, Figures 6A, 6B). In contrast to dLMPs, the early increase in the distal-posterior pLMPs populations appears transient as the spatial distribution becomes similar to wildtype forelimb buds by E11.25 (Figure 6C, 6D).

**Figure 6.**
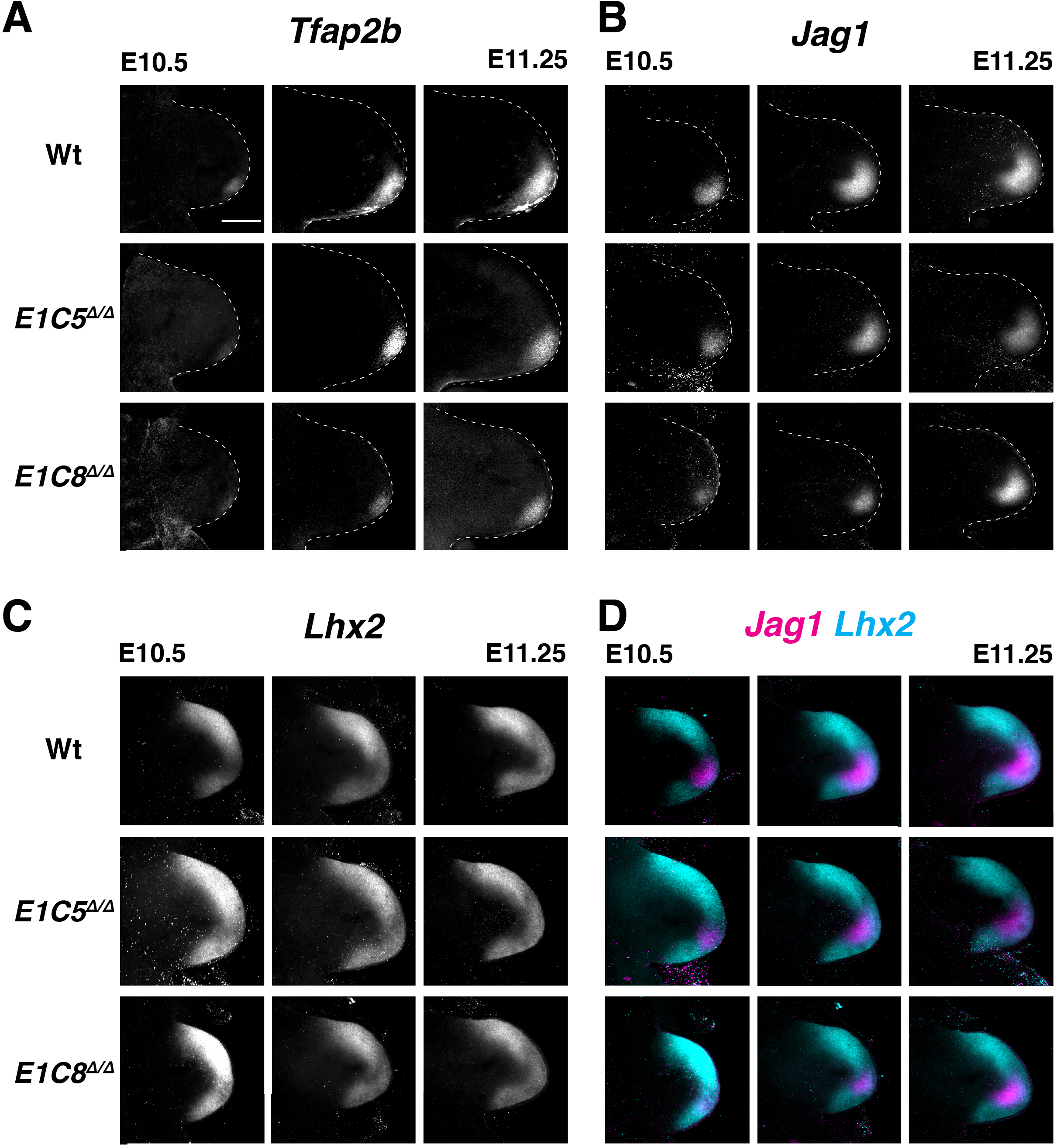
Spatio-temporal progression of the dLMP and pLMP domains. (A-D) Comparative RNA-FISH analysis of a temporally ordered series of wildtype and *E1C5*^Δ/Δ^ and *E1C8*^Δ/Δ^ forelimb buds (E10.5 to E11.25, range: 36-43 somites) illustrates the temporal progression in spatial gene expression (n≥3 replicates per genotype). All limb buds are oriented with anterior to the top and posterior to the bottom. Scale bar: 300μm. (D) Merge of the *Jag1* (magenta) and *Lhx2* (cyan) expression patterns.

### Spatial alterations in dLMP and pLMP signature gene expression foreshadows congenital digit malformation phenotypes

To gain insight into the extent to which spatial alterations in *Grem1* affect the spatial distribution of dLMP and pLMP signature gene expression, forelimb buds of mouse models for congenital limb and digit malformations were analyzed (Figures 7A-7D). In mouse forelimb buds lacking *Grem1* (*Grem1*^ΔL/ΔL^) at E10.75, the pLMP marker *Lhx2* is expressed uniformly within the distal-peripheral mesenchyme, while expression of the *Asb4* marker gene is expanded posteriorly within the distal mesenchyme in comparison to wildtype controls (Figure 7B, compare to Figure 7A). Strikingly, *Jag1* and *Tfap2b* expression are below detection, while the *Hoxa13* expression domain is spatially reduced (Figure 7B, *Hoxa13*: Figure S6A). This analysis reveals the predominant *Grem1*-requirement for *Jag1*^+^ and *Tfap2b*^+^dLMPs. Genetic inactivation of *Shh* disrupts *Grem1* expression and results in the formation of one very rudimentary digit (Figure 7C).^14,54,55^ The *Shh* deficiency alters both LMP populations, as the anterior expression bias of *Lhx2* and *Asb4* is lost, and neither *Jag1* nor *Tfap2b* expression are detected (E10.75, Figure 7C). We previously showed that in *Shh*-deficient limb buds *Hoxa13* expression is reduced to residual levels in agreement with almost complete autopod agenesis.^56^ Conversely, *Gli3* inactivation results in preaxial polydactyly and precocious anterior expansion of the *Grem1* expression domain (E10.75, Figure 7D).^56-58^ In *Gli3* ^Δ/Δ^ forelimb buds, the *Lhx2* expression levels and *Asb4* domain are reduced in particular in the anterior and distal mesenchyme (left panels, Figure 7D). In contrast, the dLMP markers *Jag1* and *Tfap2b* (right panels Figure 7B) follow the precious anterior *Grem1* expansion together with *Hoxa13* expression.^56^ This early expansion of dLMPs foreshadows the preaxial polydactyly that arises much later,^58^ while the anterior reduction in pLMPs correlates with the loss of anterior digit identities. Collectively these data indicate that changes in the dLMP population size alter digit numbers, whereas changing spatial asymmetry, i.e. losing the anterior pLMP population bias, correlates with the loss of digit identities.

**Figure 7.**
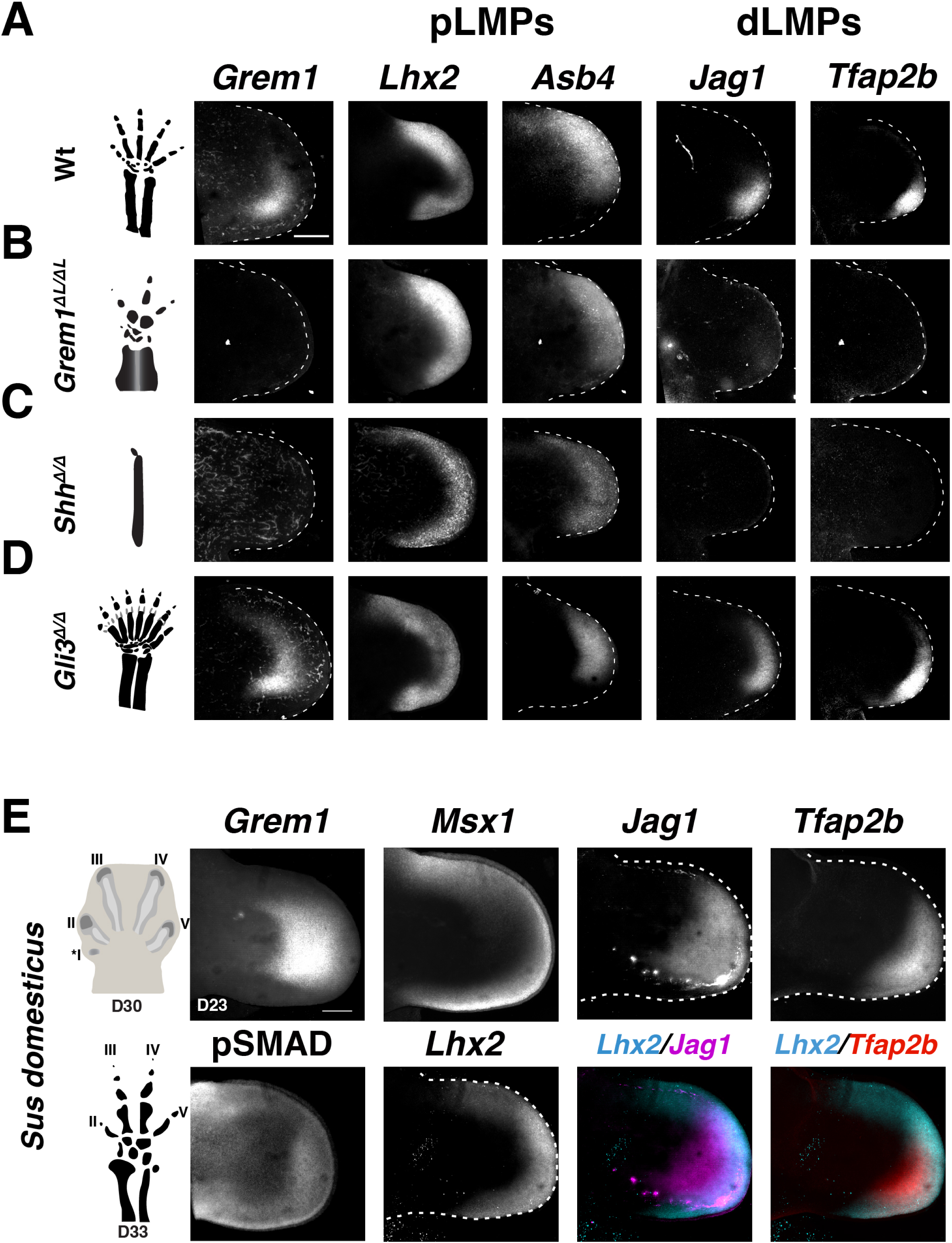
RNA-FISH analysis of *Grem1* and pLMP and dLMP marker genes in mouse models for human congenital malformations and pig limb buds. (A-D) Comparative analysis of the spatial distribution of *Grem1*; *Lhx2* and *Asb4* (pLMPs); *Jag1* and *Tfap2b* (dLMPs) in wildtype and mutant limb buds. Left-most panel: schemes of the resulting digit phenotypes at E14.5. All limb buds (E10.75) are oriented with anterior to the top and posterior to the bottom. (A) wildtype forelimb buds. (B) *Grem1*-deficient limb buds. (C) *Shh*-deficient forelimb buds (prior to the onset of major apoptosis). (D) *Gli3*-deficient limb buds. n≥3 independent biological replicates for all genotypes and probes. (E) Analysis of the spatial distribution of *Grem1*, pSMAD, *Msx1* and signature genes for pLMPs (*Lhx2*) and dLMP (*Jag1*, *Tfap2b*) pig forelimb buds at D23 (n=3 independent biological replicates per probe). Schemes (left panel) show the pentadactyl digit anlage in a pig forelimb bud at D30 and the resulting tetradactyl digit pattern at D33 (not drawn to scale, adapted from ref.^61^. For whole mount RNA-FISH pig-specific HCR probes were used and pSMAD activity was detected by whole mount immunofluorescence. Lower right panels show the merge of the *Lhx2* with the *Jag1* or *Tfap2b* expression domains in the same forelimb bud. All limb buds are oriented with anterior to the top and posterior to the bottom. Scale bars: 300μm. Relates to Figure S6.

### Molecular similarities and differences underlie the congenital digit and identity loss in mouse limb buds and the evolutionary digit reductions in pig limb buds

The symmetrical nature of *Grem1* expression in pig limb buds^23^ prompted us to gain insight into how dLMPs and pLMPs might have been altered during evolutionary diversification of Artiodactyla from the pentadactyl archetype. Orthologous stages of pig (*Sus domesticus*) limb buds were analyzed using pig-specific HCR probes (Figure 7E). The major difference between *Grem1* tetradactyl mouse and pig limb buds is that pig limb buds are still pentadactyl at gestational day D30 (schemes, Figure 7E, orthologous mouse embryonic stage E13.25)^59^, whereas the loss of middle digit asymmetry and paraxonic axis is already apparent at D24 (Figure S6B, orthologous mouse embryonic stage E12.0)^59-61^. Only during digit growth is the vestigial anterior digit I lost due to precocious restriction of AER-FGF signaling.^61^. In early pig forelimb buds (D21), *Grem1* is expressed posteriorly comparable to wildtype mouse forelimb buds at E10.5, while pSMAD activity is rather diffuse (Figure S6C, compare to Figure 1C). The m^+^LMP (*Msx1*) and pLMP (*Lhx2*) populations expand into the posterior peripheral mesenchyme, while the dLMP expression domains of *Hoxa13*, *Jag1* and *Tfap2b* are nested in the distal-posterior mesenchyme (D21: Figure S6D-E). In pig forelimb buds at D23, *Grem1* expression is expanded anteriorly, which results in the reduction of pSMAD activity in the anterior limb bud mesenchyme (Figure 7E). This is paralleled by the anterior reduction of the *Msx1* and *Lhx2* signature genes (Figure 7E), which results in loss of the anterior bias observed in mouse wild type limb buds at E10.75 (Figure 7A). In contrast, the dLMP signature genes *Hoxa13*, *Jag1* and *Tfap2b* are expressed in well-defined and spatially nested posterior-distal domains that scale with the enlarged GREM1 domain (Figure 7E and Figure S6F). Taken together, the analysis of pig limb bud development shows that, as in mouse limb buds, the dLMP population size depends on GREM1-mediated reduction of BMP activity. Similarly, the loss of the posterior *Grem1* expression bias in both mouse tetradactyl and pig limb buds results in the reduction/loss of the anterior expression bias of *Lhx2* marking the pLMP population. Thus, the loss of the anterior pLMP bias from early limb bud development onwards precedes the loss of middle digit asymmetry in both species.

## Discussion

Previous studies in vertebrates indicate that spatial differences in limb bud mesenchymal *Grem1* expression are an early molecular indicator of subsequent alterations affecting the pentadactyl pattern, whether in congenital malformations or due to the evolutionary diversification of the archetype pentadactyl pattern^24,25,62,63^. Malkmus et al.^23^ showed that conserved CRM enhancers in the genomic *Grem1* landscape^64,65^ provide the spatio-temporal regulation of *Grem1* expression and pentadactyly with *cis*-regulatory robustness. Genetic inactivation of several CRM enhancers results in significant spatial reduction and loss of the posterior *Grem1* expression bias in mouse limb buds, which underlies the loss of both digit d2 and middle digit asymmetry that results in tetradactyly (this study).^23^ The observed transition from pentadactyl mesaxonic to tetradactyl paraxonic digit development is reminiscent of the digit reductions that occurred during the evolutionary diversification of Artiodactyl limbs.^23,25,59-61,66^ In *Grem1* tetradactyl mouse limb buds, the spatial reduction of GREM1 increases BMP activity and downscales the SHH/GREM1/AER-FGF feedback signaling system (this study), whose establishment during onset of limb bud development depends on rapid GREM1-mediated reduction of BMP activity and SHH signal relay.^12,14,20,67^ These studies together with the molecular alterations underlying the loss of the anterior digit d2 in *Grem1* tetradactyl limb buds (this study) render it likely that GREM1 is the relay signal for anterior digit specification downstream of direct specification of posterior digits by SHH.^3^ This is supported by the fact that *Grem1* is required to time the mesenchymal response to SHH signaling in early limb buds.^13^ Alternatively, the alterations in *Grem1* tetradactyl limb buds could impact the long-range SHH signaling that has been proposed to pattern digit d2.^6^

Indeed, the analysis of *Grem1* tetradactyl and additional mouse mutant limb buds points to an instructive role of GREM1-mediated BMP antagonism in specifying the size of the dLMP population as: 1. the spatial reduction or expansion of the *Grem1* domain directly impacts the corresponding reduction or expansion of the dLMP population size in mutant limb buds. 2. maintenance and proliferative expansion of JAG1^+^dLMPs in culture critically depends on inhibition of BMP signal transduction^68^; 3. the expression domains of *Grem1* and dLMP signature genes are proportionally enlarged in pig limb buds in comparison to mouse limb buds, showing that across the evolutionary scale, the dLMP population size changes in sync with spatial *Grem1* expression. The positive regulation of the dLMP population is likely direct due to the spatial proximity of *Grem1*-expressing and dLMP cells. However, regulation of dLMP population size could also be indirect via the SHH/GREM1/AER-FGF feedback loop as genetic inactivation of *Fgf8* together with *Fgf17* or *Fgf9* in the AER results tetradactyly, but middle digit asymmetry is maintained.^39^ Furthermore, expression of the dLMP signature gene *Hoxa13* is the broadest along the AP axis, which agrees with the *Hoxa13* lineage contributing to all skeletal elements of the autopod.^33^ In *Hoxa13*-deficient mouse limb buds, the anterior expansion of both *Grem1* and *Jag1* expression is foreshortened, which also results in tetradactyly and loss of middle digit asymmetry^69,70^. During autopod development, *Jag1* expression is restricted to the distal mesenchyme that will give rise to the digit arch.^13,71^ In contrast to *Hoxa13* and *Jag1*, *Tfap2b* expression is much more restricted and a recent study shows that the *Tfap2b* lineage contributes to digits.^72,73^ Therefore, our single cell analysis identifies triple-positive *Hoxa13*^+^*Jag1*^+^*Tfap2b*^+^ dLMPs as early GREM1-dependent digit progenitors in mouse and pig limb buds. Moreover, dLMP signature genes are also detected in autopodial cell signatures of human limb buds.^35^ Finally, the changes in dLMP population sizes correlate positively with the resulting numbers of digits formed mutant mouse limb buds with spatially altered *Grem1* expression (this study).

In contrast, the pLMP population is increased in the distal-posterior mesenchyme of *Grem1* tetradactyl limb buds from early stages onwards, which shows that the pLMP population size correlates positively with increased BMP activity. This is corroborated by the reduction of pLMPs in the anterior mesenchyme of *Gli3*-deficient limb buds as a consequence of the anterior expansion of *Grem1* expression. Indeed, several pLMP signature genes are direct transcriptional targets of SMAD4-mediated BMP signal transduction in the anterior mesenchyme of wildtype limb buds at E10.0.^20^ This anteriorly biased pLMP distribution is indeed captured by single cell analysis together with a distal-posterior gap in the pLMP domain that overlaps the region in which the dLMP population is established. Our analysis reveals two molecular alterations underlying the loss of the anteriorly pLMP distribution in early limb buds: 1. in tetradactyl mouse limb buds, the spatial reduction and lack of posterior bias in *Grem1* expression leads to an increase in pLMPs in the distal-posterior mesenchyme, leveling out the anterior bias. In *Gli3* deficient limb buds, anterior expansion of *Grem1* causes loss of pLMP asymmetry due to reduced anterior BMP activity, which in turn promotes excess LMP proliferation in the anterior mesenchyme, which underlies the preaxial digit polydactyly^58^. In both cases, this results in loss of middle digit asymmetries, which indicates that the anterior bias of the pLMP population in early limb buds is required to specify AP digit polarity, with is functionally genetically substantiated by the loss of AP polarity and digits in *Lhx2/9* deficient mouse limbs^52^.

Analysis of congenital limb malformations and evolutionary limb diversification has shown that spatial divergence from the archetype pentadactyl *Grem1* pattern is the first molecular indicator of the resulting digit pattern. For example, *Grem1* has been proposed as an early “sensor” of digit patterning in different bird embryos as the size of its expression domain correlates positively with digit numbers.^74^ Analysis of *Grem1* in species from different clades including basal fishes reveals the amazing spatial plasticity in expression that reflects *cis-*regulatory diversification^23^. For artiodactyl species, it has been proposed that early spatial alterations in *Grem1* prefigure the paraxonic nature and digit reductions in bovine and pig embryos,^23,60,61,66^, but the the underlying cellular alterations remained unknown. Pig limb buds are initially pentadactyl but exhibit paraxonic characteristics from the early stages onward.^59-61^. Molecular analysis of dLMP and pLMP signature genes in pig limb buds shows that the dLMP population scales with the increase in GREM1-mediated BMP antagonism, i.e. is not reduced as observed for the early loss of digit d2 in *Grem1* tetradactyl mouse limb buds. In contrast, the anterior bias of the pLMP population is lost in both types of tetradactyly: in pig limb buds *Lhx2*^+^pLMPs are reduced anteriorly while *Grem1* tetradactyl mouse limb buds pLMP are increased distal-posteriorly. Both genetic and evolutionary analyses lend support to the conclusion that the reduced or absent anterior bias in pLMPs is linked to the loss of middle digit asymmetry, which is a defining feature of unguligrade posture in Artiodactyla.^24,25^ Last but not least, the molecular analysis of signature genes concurs with the fossil record which established that the transition from mesaxony to a paraxony preceded digit reductions and loss^25,75^. Both these genetic and evolutionary data provide a likely and rather straight forward explanation for the observed significant plasticity of digit numbers and identities in tetrapods. The spatial regulation of *Grem1* as part of the self-regulatory signaling system^1,12,14-18^ can be likened to a control dial that tunes the balance of BMP activity (pLMPs: up; dLMP down) and BMP antagonism (pLMPs: down; dLMP up) in space and time. During tetrapod evolution and diversification of the archetype pentadactyl pattern, shifting the balance of this integrative system in either direction would have impacted one or both LMP populations in ways similar, but not identical, to those seen in various congenital digit malformations. The vast diversity in tetrapod digit patterns even among rather closely related species such as primates,^76^ Artiodactyla,^25^ squamates and birds^74,77^ points to the existence of such a tunable control dial that provides an otherwise robust regulatory system with evolutionary plasticity.^23^

## Limitations of the study

Comparative single-cell RNA sequencing identifies two early specified LMP populations with characteristic molecular signatures that both participate in digit patterning. While they provide a cellular blueprint to analyze the specification of digit numbers and identities in different species and alterations caused by congenital malformations, deviations from the pentadactyl ground state that result in digit loss and reductions can also occur later, likely involving others mechanisms.

## Acknowledgements

The authors would like to thank A. Offinger and her team for outstanding mouse husbandry. Additionally, we would like to thank the P. Pelczar from the Center for Transgenic Models (CTM), University of Basel and his team for the generation genome-edited founder mice. We are grateful to N. Tichy for the whole mount *in situ* hybridization analysis shown in Figure S2. We would like to thank P. Lorentz from the DBM Microscopy Core Facility for technical support throughout the project and L. Sauteur for providing support for image processing. In addition, we are grateful to the Biozentrum Imaging Core Facility for support with light sheet fluorescence microscopy, especially to S. Roig, K. Schleicher and L. Gerard. We thank B. Yoder for providing the *Alx4*-Cre^ERT2^ mouse. We are grateful to T. Aguirre-Lavin for providing us with pregnant sows and the lab space to collect embryos. This research was supported by grants from the ERC advanced grant INTEGRAL ERC-2015-AdG; Project ID 695032 (to R.Z.), the Swiss National Science Foundation (SNSF): 310030_166685B to RZ and AZ, 310030_184734 and 310030_207824 to RZ with AZ as project partner, 310030_192604 to B.T., the National Center of Competence in Research Molecular Systems Engineering to B.T. and the University of Basel provided core funding (to A.Z. and R.Z.).

## Author contributions

A.Z. and R.Z. conceived and supervised the study. Figures were prepared and the manuscript was written by V.P., A.P., A.M., R.Z. and A.Z. with input of all authors. J.M. generated the *Grem1*^ΔE1C8^ mouse mutant. A.P. performed the lineage analysis. A.M. and A.P. performed the immunostainings. V.P. performed the single-cell experiments; generated and curated the datasets and performed the bioinformatics analysis with advice and input from Z.H. and B.T.; interpretated the single-cell data together with A.Z., V.P., A.M. and A.P. performed the RNA-FISH analysis with contributions from G.S. A.Z. and R.Z. collected pig embryos, and A.M. performed the analysis of pig limb bud. All authors discussed the results and commented on the manuscript.

## Methods

### EXPERIMENTAL MODELS

### Animals

#### Ethics statement and approval of all animal experimentation

All animal studies present in this manuscript were performed in accordance with national laws and approved by the national and local regulatory authorities as mandated by law in Switzerland and France. Mouse studies were approved by the Regional Commission on Animal Experimentation and the Cantonal Veterinary Office of Basel (national license 1950) in accordance with Swiss laws and the 3R principles.

#### Mouse strains and embryos

In line with the refine and reduce 3R principles, all strains were bred into a Swiss Albino (*Mus musculus*) background as only robust phenotypes manifest in this strain background and the numbers of embryos and litter sizes are large (≥12-15 embryos per pregnant females). Embryos of both sexes at the developmental ages indicated were used for experimental analysis. Embryos were age-matched by counting somites and matching limb shapes and sizes. The following genetically modified mouse strains were used in this study: two alleles with spatially restricted and reduced *Grem1* expression due to deletion of CRM enhancers in the *Grem1* landscape, namely *EC1CRM5*^Δ/Δ^ (*E1C5*^Δ/Δ^), ^23^ and *EC1CRM8*^Δ/Δ^ (*E1C8*^Δ/Δ^, this study); for lineage analysis the *Shh*GFPCre (inserted into the *Shh* gene),^6^ and *Alx4*-Cre^ERT2^ (random transgene insertion),^42^ mouse strains were used in combination with the *ROSA26* ^LSL−*tdTomato*41^ and *ROSA26R*−GFP^43^ reporter mice respectively. *Shh*-deficient embryos were generated using the *ShhGFPCre* strain and *Gli3*-deficient embryos using the *Gli3^Δ^* strain.^58^

#### Generation of the *EC1CRM8*^Δ/Δ^ (*E1C8*^Δ/Δ^) *Grem1* allele

CRM8 is highly conserved 736bp genomic element (present in cartilaginous fish)in open chromatin but LacZ reporter analysis in mouse limb buds did not reveal obvious enhancer activity (Malkmus, et al. 2021). Its ancient origin and distal genomic position in proximity to the 3’border and CTCFs sites within the *Grem1* TAD prompted functional analysis in the context of the *EC1* deficiency by genome editing. Two single guide (sg)RNAs were designed to target the 2596 bp region of open chromatin that includes the highly conserved core region of CRM8. The sgRNAs and Cas9 protein were delivered by electroporation to *EC1*^Δ/Δ^ zygotes. Founders were genotyped for the *CRM8* deletion and *EC1*^Δ^ allele. The accuracy of the *CRM8* deletion was verified by sequencing the breakpoint regions. After breeding founders to the Swiss Albino mice, the deletion of *CRM8* in *cis* to *EC1* was reconfirmed by PCR analysis and sequencing. Initial analysis of *EC1CRM8* ^Δ/Δ^ (*E1C8*^Δ/Δ^) revealed the tetradactyl limb phenotype and spatial reduction of *Grem1* expression (Figure 1A-C), which resulted in inclusion of the *E1C8*^Δ/Δ^ allele in this study.

#### Pig embryos

Pig (Sus domesticus) embryos were obtained from artificially inseminated sows destined for meat production. Embryos were collected at the relevant orthologous stages (D21-D24)^61^ at the facility of INRAE Val de Loire.

### Method details

#### Skeletal analysis

For limb skeletal preparations, embryos were collected on day 14.0 and embryonic stages assigned using ossification (E14.0-E14.75). Briefly, embryos were collected in PBS and fixed in 95% ethanol (Carl Roth, T171.7) overnight. After staining for 24hrs in 0.03% (w/v) Alcian blue (Sigma-Aldrich, A3157), 80% ethanol, 20% glacial acetic acid (Sigma-Aldrich, 100063) they were washed for 24 hrs in 95% ethanol. Next, embryos were pre-cleared for 30min in 1% KOH (Sigma-Aldrich, 105033) and counterstained in 0.005% (w/v) Alizarin (Sigma-Aldrich, A5533) in 1% (w/v) KOH. Finally, embryos were cleared in stepwise increased concentrations of glycerol/1% KOH (20%,40%, 60%, 80% glycerol) and stored in 80% glycerol in water. Alcian blue detects cartilage and alizarin red ossified bone. n≥3 embryos were analyzed per genotype.

#### Fluorescent whole mount HCR^TM^ RNA *in situ* hybridization (RNA-FISH)

The mouse HCR^TM^ probes for the different mouse genes analyzed were purchased from Molecular Instruments (USA). Briefly, embryos were fixed in freshly prepared 4% paraformaldehyde (Sigma-Aldrich, P6148) overnight at 4°C and dehydrated into 100% methanol (VWR, 20903.368) for storage at -20°C. The RNA-FISH analysis and image acquisition was done exactly as described in the step-by step protocol.^36^ For some images acquired by confocal imaging (see below) autofluorescence was removed by image processing. This was done using FIJI to display the maximum projection of the DAPI channel, or sub-stacking to exclude tissue below or above the limb bud (i.e. remnants from dissection and embedding) by generating a selection outside the limb bud. Briefly, this projection was thresholded using the mean threshold method. Holes were filled in the binary image and if necessary, iterative binary close or open function was applied to clean the mask. A region outside the limb bud was selected and applied to the original multi-channel stack and pixel values for each channel and slice were set to 0. The selection was saved. Finally, brightness and contrast were adjusted for each channel. ImageJ macroscripts are available in a Zenodo repository.

#### Whole mount immunofluorescence

All embryos were dissected in ice-cold PBS and fixed subsequently in 4% PFA overnight. After three 5 min washes in PBS embryos were transferred to 0.1%Tween (Sigma-Aldrich, 93773) in PBS (PBS-T) for splitting embryos in half and stored in 0.01% sodium azide (Sigma-Aldrich, S2002) in PBS at 4°C. For detection of GREM1 and phospho-SMAD1.5.9 (pSMAD1.5.9) proteins, samples were progressively dehydrated to 100% Methanol in 0.01% PBS-T and subsequent steps were performed as described in ref.^36^ with the following modifications. Incubations with primary antibodies were performed in primary antibody solution: 1%BSA (Sigma-Aldrich, A2153), 10% donkey serum (Sigma-Aldrich, S30), and 0.5% Triton (Sigma-Aldrich, T8787) in PBS-T for 48-72 hours with gentle shaking at 4°C. Anterior lineage analysis used as primary antibodies rabbit-anti-SOX9 (1:400, Millipore, ab-5535) and sheep-anti-GFP (1:400, Bio-Rad, 47451051). For posterior lineage analysis rabbit-anti-dsRed (Takara, 632496) and goat-anti-SOX9 (R&D, AF3075) primary antibodies were applied. For GREM1 and pSMAD proteins: primary antibody incubations were performed in primary antibody solution but with 0.4% Triton in PBS. For detecting GREM1 we utilized (1:100, R&D Systems, AF956) and for p-SMAD1.5.9 we applied (1:100, Cell Signaling Technology, 13820S) as primary antibodies. For the detection of phospho-ERK1/2 (pERK) and SOX9 the dehydration and bleaching steps were omitted. Primary antibodies used to detect the pERK proteins were rabbit-anti-pERK (1:200, Cell Signalling Technology, 9101) and goat-anti-SOX9 (1:400, R&D, AF3075) in primary antibody solution. Following primary antibody incubations, all samples were washed for 6x30 min with 0.5%Triton in PBS at room temperature and incubated with secondary antibodies in secondary antibody solution (1%BSA, 10% donkey serum and 0.5% Triton in PBS) for 48-72 hrs with gentle shaking at 4°C. For anterior lineage analysis donkey anti-rabbit-555 (Invitrogen, A31572) or donkey anti-rabbit 647 (Invitrogen, A31573) and donkey anti sheep-488 (Jackson, 713-545-147) antibodies were used. Posterior lineage analysis utilized donkey anti-rabbit-488 (1: 250, Jackson,711-545-152), donkey anti-goat-555 (1:250, A-21432) or donkey anti-goat 647 (1:250, Invitrogen, A-21447). For GREM1 protein and pSMAD1.5.9. protein the same secondary antibody solution was used with 0.4%Triton in PBS and an incubation of 24h with the following secondary antibodies donkey anti-goat 647 (1:1000, Invitrogen, A21447) and donkey anti-rabbit 555 (1:1000, Invitrogen, A31572). For pERK and SOX9 immunofluorescence donkey anti-rabbit 647 (1:250, Invitrogen, A31573) and donkey anti-goat-555 (1:250, Invitrogen, A-21432) were applied and were incubated between 48-72 hours in secondary antibody solution. For all samples cell nuclei was counterstained with DAPI (Sigma-Aldrich, D9542) (1:1000) diluted into the secondary antibody solution before starting the sample clearing.

##### Hydrophilic and hydrophobic clearing of samples for cell lineage analysis

Hydrophilic samples were cleared in 2.5M Fructose-Glycerol 60% (v/v) (Fructose: Sigma-Aldrich, F0127; Glycerol: AppliChem, 131339) and mounted as described.^36^ Alternatively, hydrophobic tissue clearing was performed as described <u>here</u>. Briefly, forelimb buds with flanks were embedded in 2% Agarose (Promega, V3121) prepared in distilled sterile water. Agarose cubes containing the samples were dehydrated through a graded methanol series starting with 20% ending with 100%, 30min-1hr each and an additional step in 100% methanol for 1hr. Samples were placed into 1/3 Methanol and 2/3 dichloromethane (DCM, Sigma-Aldrich, 270997) with rotation at 13rpm at room temperature for overnight incubation. The following day, the mixture was replaced with 100% DCM for 30 min with rotation at 13rpm at room temperature. Subsequently, the 100% DCM solution was removed and dibenzyl ether (DBE, Sigma-Aldrich, 33630) was added by filling the tube to the top to prevent oxidation. Samples were stored in DBE protected from light until imaging. Imaging was performed after refractive index (RI) equilibration for at least 2 hours prior to imaging in ethyl cinnamate (ECI, Sigma, 112372) and then placed again in DBE to preserve fluorescence following image acquisition.

#### Fluorescent image acquisition and processing

*Confocal spinning disk microscope acquisition:* after hydrophilic tissue clearing, limb buds were imaged using a 10x objective (10x/0.45 CFI Plan Apo) with a confocal spinning disc scan unit (Yokogawa Spinning Disk CSU-W1-T2) and a Nikon Ti-E, Hamamtsu Flash 4.0 V2 CMOS camera. The image acquisition software VisiView Premier was used to set the acquisition parameters. Z-step size was set to 5 µm for the lineage analysis and to 2.3 µm for RNA-FISH to generate the 2-dimensional maximum projection. The raw *nd* file was converted to an *ims* file using the IMARIS file converter (9.9.1) and by placing the correct voxel sizes. If necessary auto-fluorescent blood vessel were removed as described below.

*Light-sheet fluorescence microscope acquisition*: after hydrophobic tissue clearing, Zeiss Lightsheet 7 microscope with the ZEN black 3.1 LS (version 9.3.10.393) software and lasers at a fixed wavelength of 405 nm, 488 nm, 561 nm and 638 nm was used. Dual side illumination was performed using the illumination air lenses LSFM foc 10x/0.2 and adjusted for an RI of 1.55. Fluorescence was detected using an Clr Plan-Neofluar 20X/1.0 immersion detection objective, with the correction collar adjusted to the refractive index. Agarose-embedded samples were adhered to a metal holder and mounted onto the sample holder. Then, the samples were immersed into a 20x clearing chamber filled with 35–45 mL of ECI. Manual alignment of the light sheet was performed using the 561 nm laser and adjusting the collars to the refractive index. The Z-stacks were acquired with the full pco.edge cameras chip (1920X1920 pixels), zoom 0.36, using the optimal step size of 1.27 µm to achieve the recommended Nyquist sampling. All acquired images were aligned and fused with Fiji and custom scripts (https://doi.org/10.5281/zenodo.8019804) using the BigStitcher plugin (https://doi.org/10.1038/nmeth0610-418). The results were then converted to an IMARIS file that allows the reconstruction of the 3D volumes using Imaris (Bitplane, 9.9.1). If necessary auto-fluorescent blood vessels were removed in IMARIS by generating a surface using the 647 channel for autofluorescence (icx) and by masking the generated blood vessel surface in the 488 channel.

#### Image processing for cell lineage analysis

The section view in Imaris was used to visualize the data along the three coordinate axes using the mean intensity projection. Section thickness and location was defined via setting in the xy view from digit d1 to digit d3 in the wildtype and from digit d1 to digit d4 in *Grem1* tetradactyl limb buds (anterior lineage) and from digit d5 to digit d3 in the wildtype and digit d5 to digit d3* in mutant limb buds (posterior lineage). Section thickness was set from the yz view and covered the whole thickness of the SOX9 digit domain. RGB TIF files were generated with Imaris and the two section views xy, and yz were used. OMERO [Software] https://www.openmicroscopy.org/omero/) was used to generate the insets of these two views. Virtual merge of anterior and posterior lineages in limb buds for better visualization of the domains. For this, age- and shape-matched forelimb buds of anterior and posterior lineage samples acquired by confocal microscopy were used. We employed Fiji (version App 2023-02-172.14.0/1.4f)^78^ to create anterior and posterior maximum intensity projections of matched forelimb buds using the DAPI channel and scaled and orientated them to match well. The larger image was scaled down to fit the smaller one (in number of image pixels). The posterior DAPI channel was aligned with the anterior one using the Fiji plugin MultiStackRegistration^79^ in combination with scaled rotation transformation. The generated transformation file was then used to align the two channels of interest for the anterior and posterior lineages. Finally, the transformed anterior and posterior channels were merged and brightness and contrast adjusted. An ImageJ macro script is available in the Zenodo repository.

#### Differential gene expression analysis using single cell RNA-seq analysis

##### Processing of limb buds for scRNA-seq

Mice were set for timed matings and embryos were collected in ice cold PBS at E10.75 (37-39 somites). Pairs of forelimb buds were dissected from selected embryos and placed into Eppendorf tubes (n= 3 biologically independent replicates per genotype). For enzymatic digestion, PBS was removed and exchanged for Collagenase D (Sigma-Aldrich, 11088858001) (20 mg/ml) in DMEM (Thermo-Fisher, 41966-029) and incubated in a water bath at 37°C for 20min. Limb buds were pipetted gently using a P1000 every 5min to help dissociation. Enzymatic digestion was stopped by adding 500 μl of HBSS+ (HBSS, 5% Fetal Bovine Serum, 1% HEPES (Thermo Fisher, 15630-080) 1M, 1% Pen-Strep (Thermo Fisher, 15140-122)) and the cell suspension was filtered and transferred to a new tube using a FlowMi Cell Strainer (Sigma-Aldrich, BAH13600040) (40μm) in combination with a P1000 tip. Tubes were centrifuged for 5 minutes at 2000g (4°C). Cells were resuspended in 100 μl HBSS+ and cell viability and numbers were assessed using a Trypan Blue (Thermo Fisher,15250061) assay on the cell counter. Libraries were generated immediately by the University of Basel Life science genomics facility. For n = 2 of Wildtype, *E1C5*^Δ/Δ^ and *E1C8*^Δ/Δ^, libraries were prepared in the lab, using Single cell library preparation system Chromium X (10X Genomics) and using the kit and manufacturer’s guidelines from the user guide Chromium Next GEM Single Cell 3’ Reagent Kits v3.1. Verified Libraries were pooled and sequenced using NovaSeq 6000 from Illumina.

#### Analysis of the primary sequencing data

We used CellRanger (v.6.0.1) with the default parameters to obtain transcript count matrices aligning the sequenced reads to the mouse genome and annotations. Count matrices were further processed using the R package Seurat (v.4.3.0.1). In the first instance, cells were filtered on the basis of the number of features expressed, number of counts and percentage of mitochondrial genes expressed. The threshold for features was > 2500, for reads was < 200000 and for mitochondrial fraction was < 0.1. All samples were merged into 1 object (merge()) and counts were normalized (NormalizeData). Next, the cell cycle was assessed using the Seurat function CellCycleScoring. Following, counts were *z*-scaled regressing out cell cycle and orig.ident (ScaleData). Principal Component Analysis (Seuratfunction RunPCA) was performed in order to obtain a two-dimensional representation of the data, RunUMAP was performed for dimensions 1:30 and otherwise with default parameters. To cluster cells, the functions FindNeighbors and FindClusters were used, the later with a resolution of 0.8. FindAllMarkers was applied to the object to find specific markers for the previously generated clusters. Based on the markers, data were subset to keep only the desired clusters of mesenchymal cells. Data were re-clustered following the same procedure. Module scores for positional and progenitor populations were determined using the function AddModuleScore and the UMAPs show color only for positive values, with a scale ranging from minimum to highest score. UMAPs for single genes expression were done with the option order = TRUE unless stated otherwise. When expression of 2 genes is in the same UMAP, the option used was blend = TRUE, with blend.threshold = 0.1 and order = TRUE.

For the Differential Expression (DE) analysis between mutants and wildtype cells, the Wilcoxon’s rank sum test was applied. Tests were executed pairwise between wildtype versus *E1C5*^Δ/Δ^ and wildtype versus *E1C8*^Δ/Δ^. Differentially expressed genes from both tests were intersected and the shared ones were highlighted in the Volcano plots shown in Figure 4 and Figure S4. X and Y chromosomal, mitochondrial and ribosomal genes, and genes expressed in blood were removed from the plots. To determine the percentage of cells in the different populations shown in barplots, different criteria were used combining single cell expression, HCR visualization and literature. For values of m^+^LMPs, we considered cells expressing *Msx1* > 0.5. pLMP cells are *Lhx2* > 0.5 | *Msx2* > 0.25 | *Rspo4* > 0.25 & *Asb4* > 1 & *Sox9* < 1. For dLMPs, *Jag1* > 0.3 | *Tfap2b* > 0.3 | *Hoxa13* > 0.3 & *Sox9* < 1.

### STATISTICAL ANALYSIS

For the Differential Expression (DE) analysis between mutants and wildtype cells, the Wilcoxon’s rank sum test was applied. Tests were executed pairwise between wildtype versus *E1C5*^Δ/Δ^ and wildtype versus *E1C8*^Δ/Δ^ limb buds.

## Supplemental Information

### Supplemental Figures

**Figure S1.**
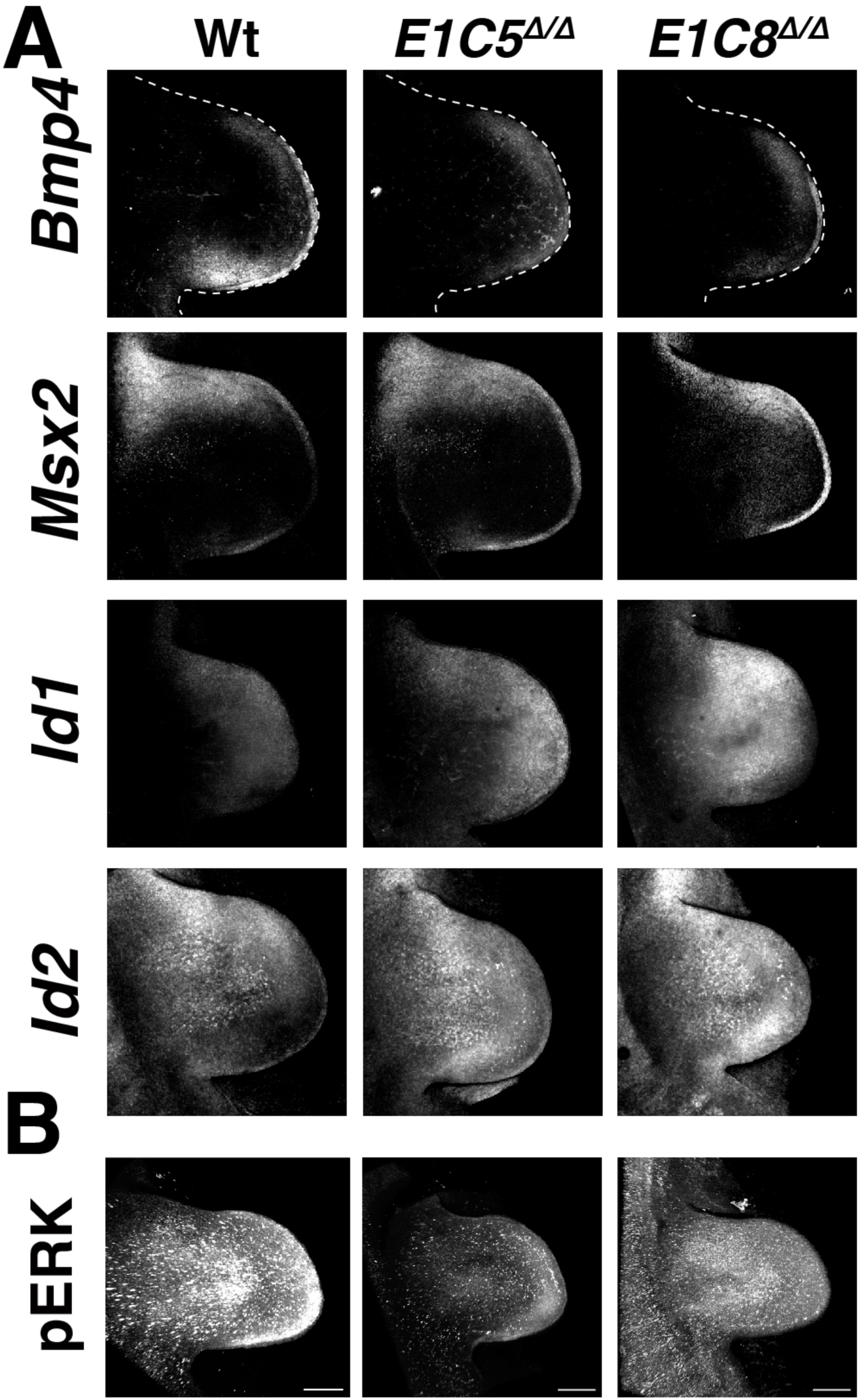
Alterations in BMP and FGF pathway activities. Comparative RNA-FISH analysis of wildtype and mutant forelimb buds at E10.75 (37-39 somites). Greyscale are shown for all analysis. (A) Analysis of BMP4 ligand in the feedback signaling system and the downstream BMP activity sensors *Msx2*, *Id1* and *Id2*. Scale bar: 300 μm. (B) Whole mount immunofluorescence analysis of phospho-ERK (pERK) activity in E10.75 forelimb buds (n=2/2 independent biological replicates per genotype). Scale bar: 200μm. All limb buds are oriented with anterior to the top and posterior to the bottom.

**Figure S2.**
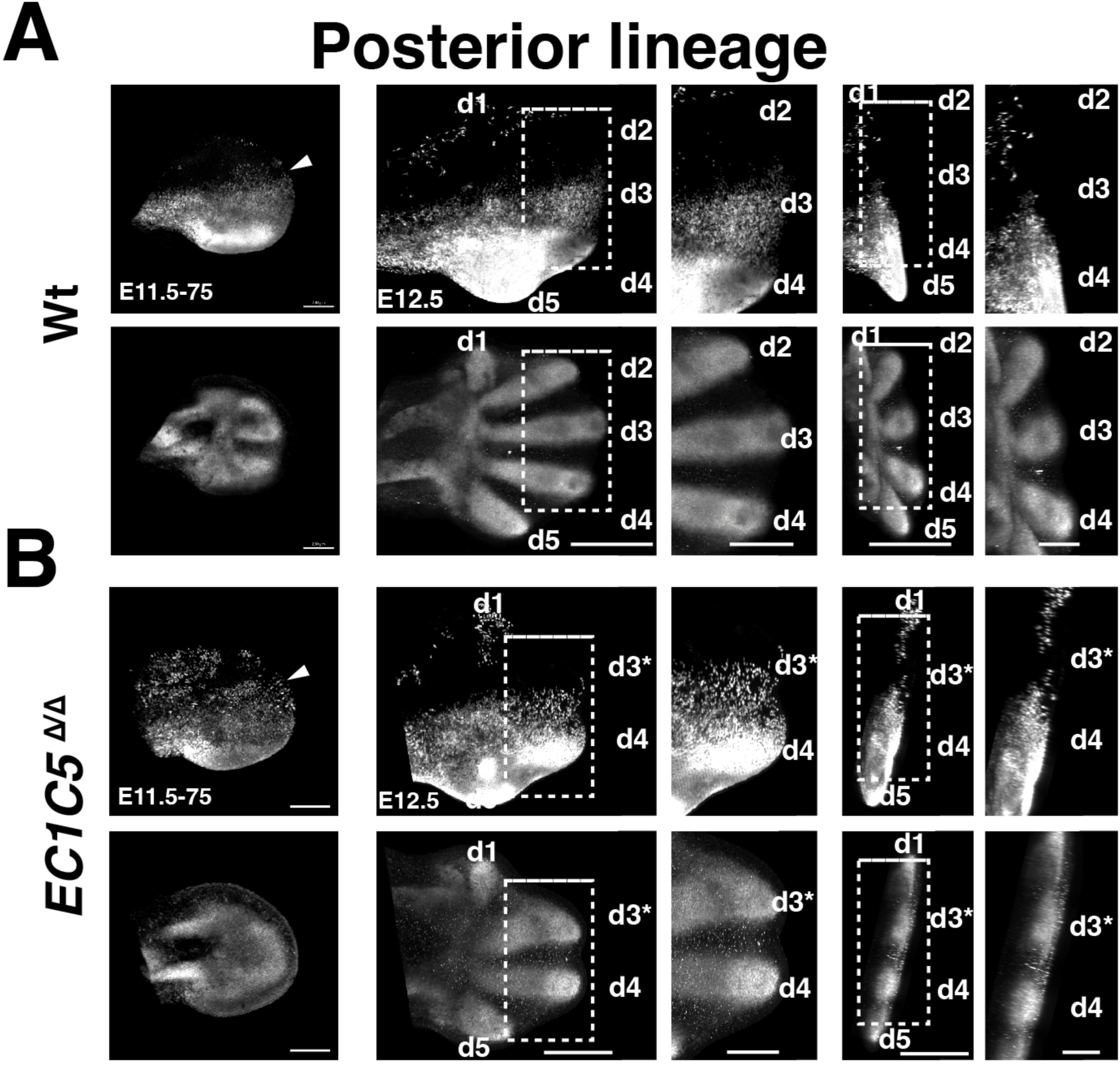
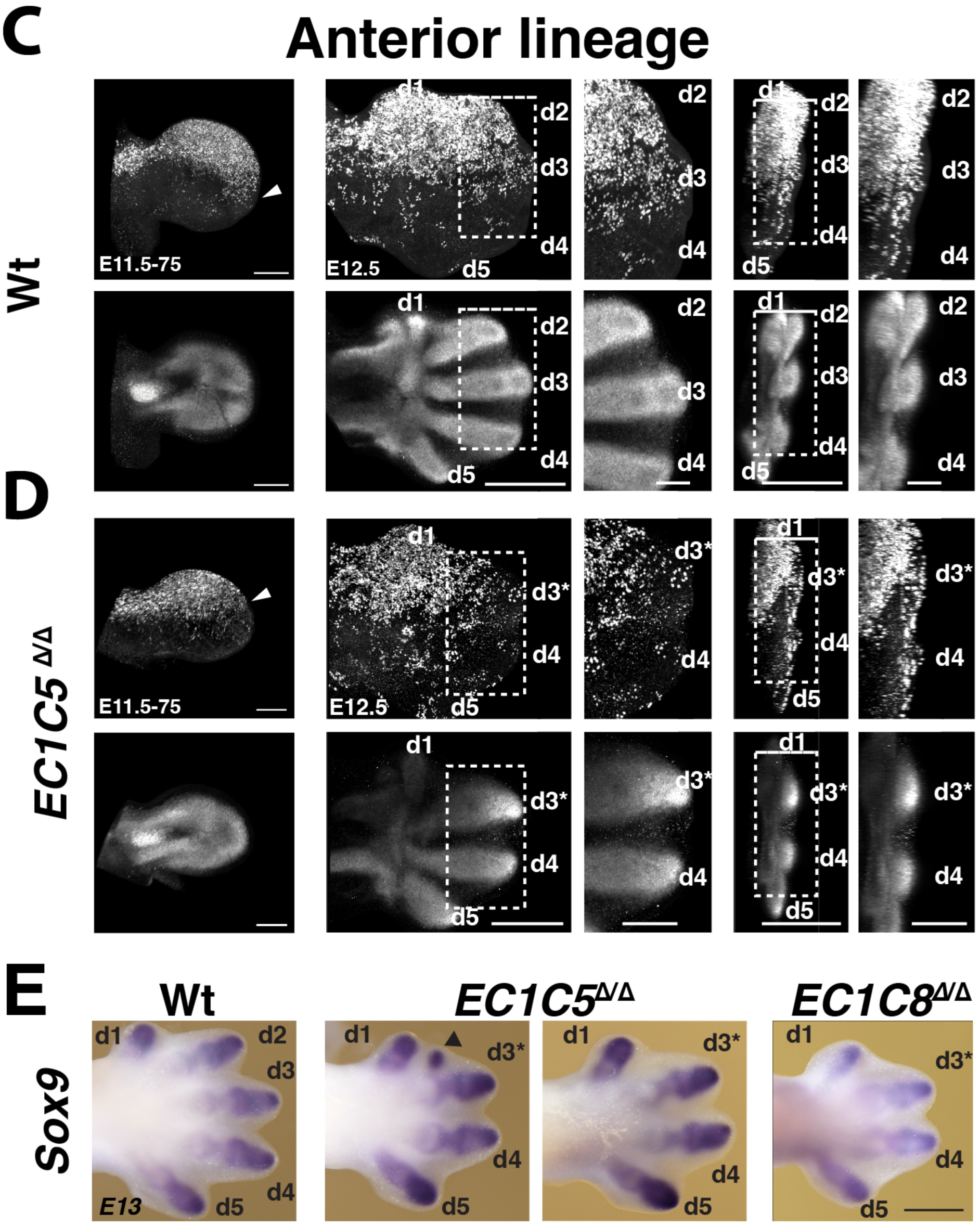
Greyscale for the lineage analysis (Figure 2) and detection of *Sox9* expression in the most anterior interdigit of some *E1C5*^Δ/Δ^ forelimb buds. The greyscale images for all posterior and anterior lineage analysis in forelimb buds shown in Figure 2 are included here. (A-B) Posterior lineage in wildtype and tetradactyl *E1C5*^Δ/Δ^ forelimb bud forelimb buds shows the posterior lineage (upper panels) and *Sox9* domain in the developing cartilage primordia (lower panels). Arrowheads indicate the anterior boundary of the posterior lineage. (C-D) Anterior lineage analysis in the wild-type and *E1C5*^Δ/Δ^ tetradactyl limb buds. Arrowheads indicate the posterior boundary of the anterior lineage. Scale bars for panels A-D: left panels: 200µm; right panels: 500 µm for overviews, 250 µm for insets. (E) Conventional whole mount RNA in situ hybridization shows the *Sox9* expression pattern in wildtype and *E1C5*^Δ/Δ^ and *E1C8*^Δ/Δ^ tetradactyl forelimbs at E13.5. Arrowhead points to the *Sox9-*positive condensation in the interdigit between digit d1 and d3* in a *E1C5*^Δ/Δ^ forelimb (n=2/5). No ectopic *Sox9*-positive condensations were detected in *E1C8*^Δ/Δ^ forelimbs (n=0/3). Scale bar: 250µm. All limb buds are oriented with anterior to the top and posterior to the bottom.

**Figure S3.**
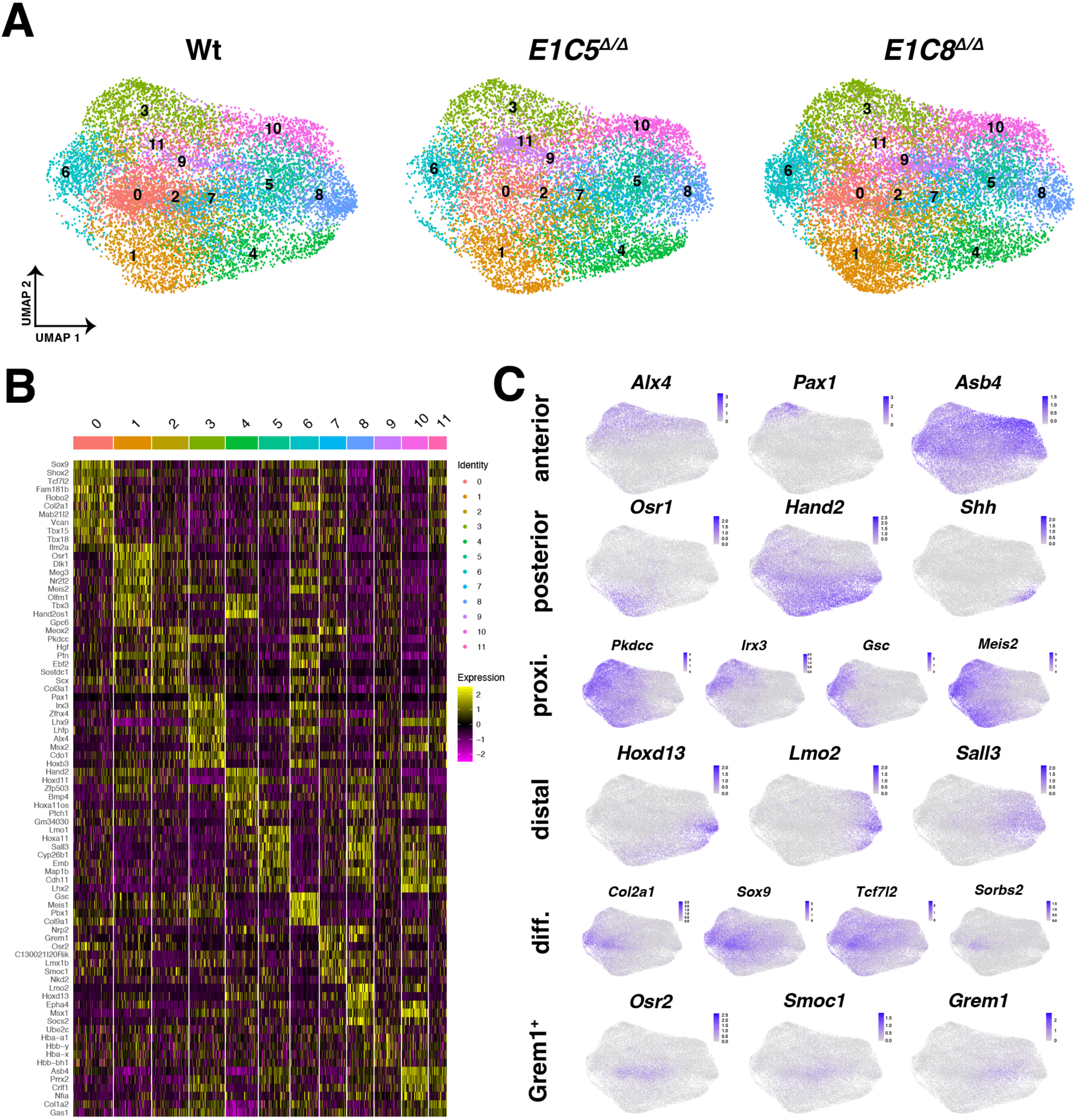
Comparative scRNA-seq analysis of wildtype and tetradactyl mouse forelimb buds (E10.75). (A) UMAP embedding of the individual scRNA-seq data for all three genotypes. (B) The pooled scRNA-seq datasets from all three genotypes were used to generate a heatmap showing the top-10 DEGs for all 12 clusters. (C) UMAP embedding of the pooled scRNA-seq datasets from all three genotypes for the genes selected to define the scores for the different limb bud mesenchymal regions. For all UMAPs, cells were plotted in random order: FeaturePlot (object, order = F).

**Figure S4.**
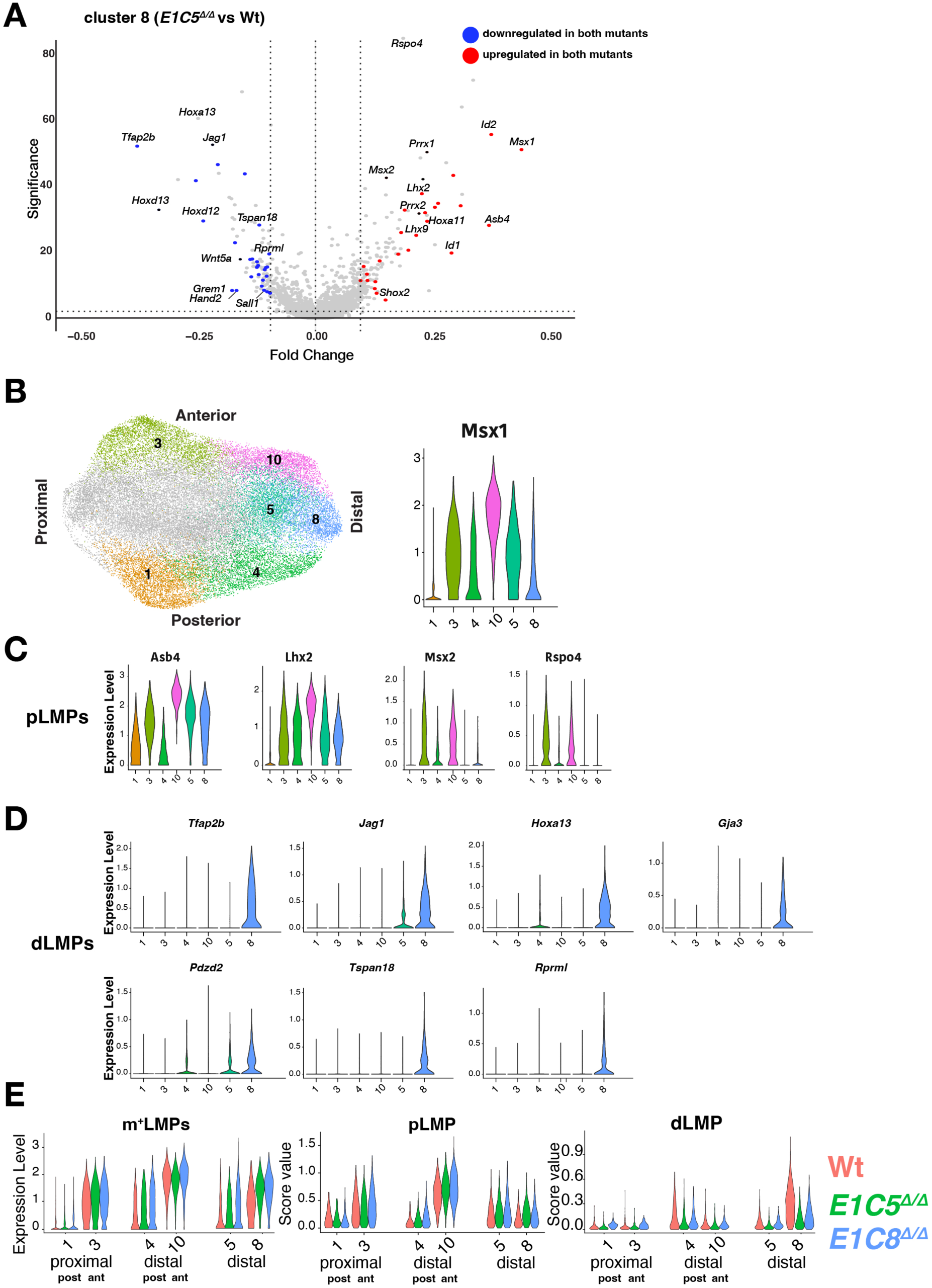
Definition of LMP genes signatures and their expression in different clusters. (A)Volcano plot shows the differentially expressed genes (DEGs) in cluster 8 by comparing *E1C5* ^Δ/Δ^ to wildtype limb buds. DEGs indicated in blue are downregulated in *Grem1* tetradactyl limb buds in comparison to wildtype limb buds. Conversely DEGs shown in red are upregulated in *Grem1* tetradactyl limb buds. Fold changes (x-axis) are shown as Log2 values. Significance (y-axis) is Log2 of the adjusted p-values. (B) Left panel: UMAP shows the relevant peripheral and distal clusters. Right panel: violin plot shows the relative expression levels of *Msx1* in all clusters shown in the left panel. (C,D) Violin plots showing the relative expression levels for the signature genes used to define pLMPs (panel C) and dLMPs (panel D) in wildtype limb buds. Their expression in peripheral and distal clusters is shown. (E) Violin plots for m^+^LPMs (*Msx1* expression levels), pLMPs and dLMPs (score values for signature genes). Wildtype values and distributions are compared with *E1C5* ^Δ/Δ^ and *E1C8* ^Δ/Δ^ forelimb buds (E10.75).

**Figure S5.**
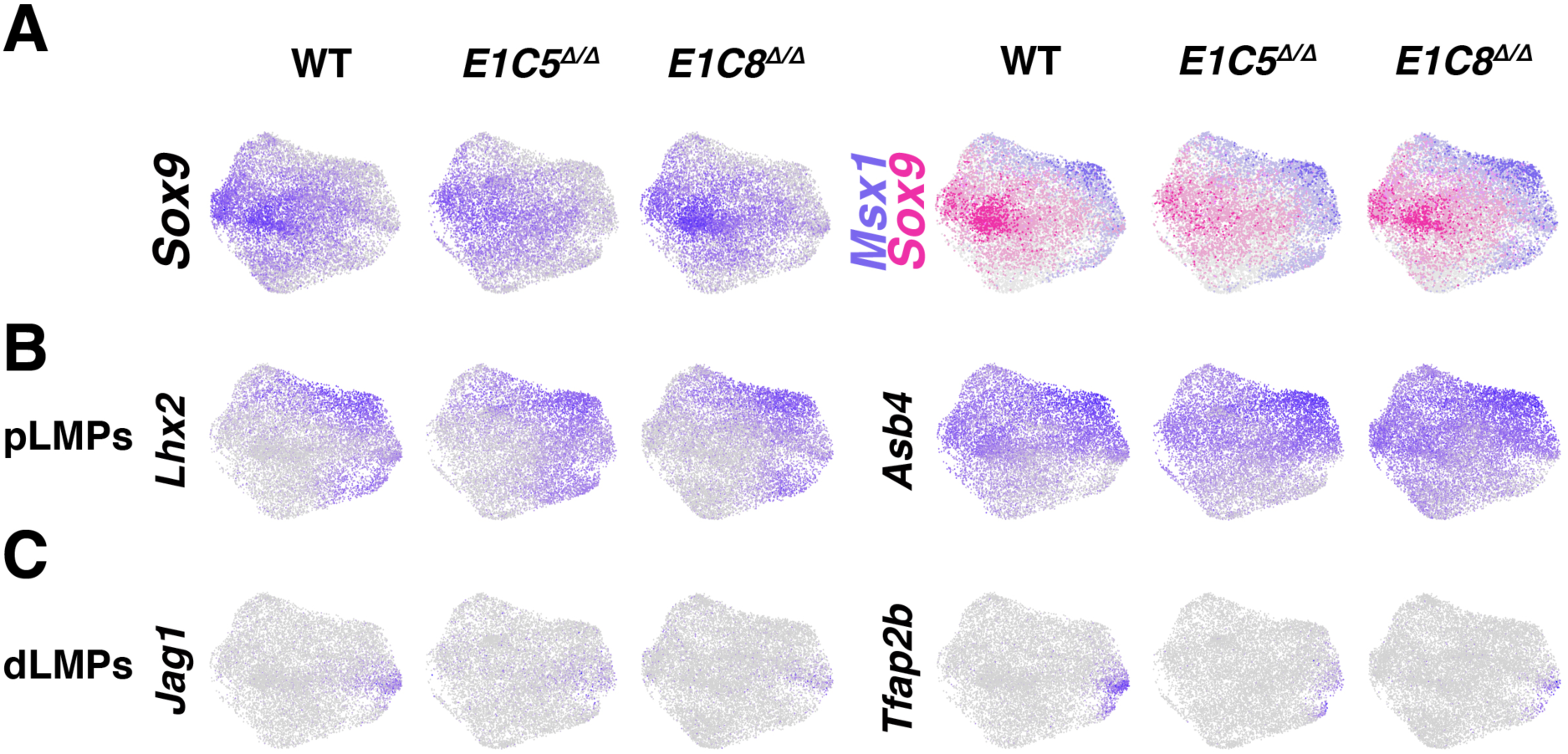
UMAP embedding of individual marker genes reveals the spatial distribution and mixing of pLMPs with dLMPs in *Grem1* tetradactyl forelimb buds. (A) UMAP embedding of *Sox9* (OCPs) and *Msx1* (m^+^LMPs) shows their complementary distribution. (B) UMAP embedding of *Lhx2* and *Asb4* shows the pLMP distribution. (C) UMAP embedding of *Jag1* and *Tafap2b* shows the dLMP distribution.

**Figure S6.**
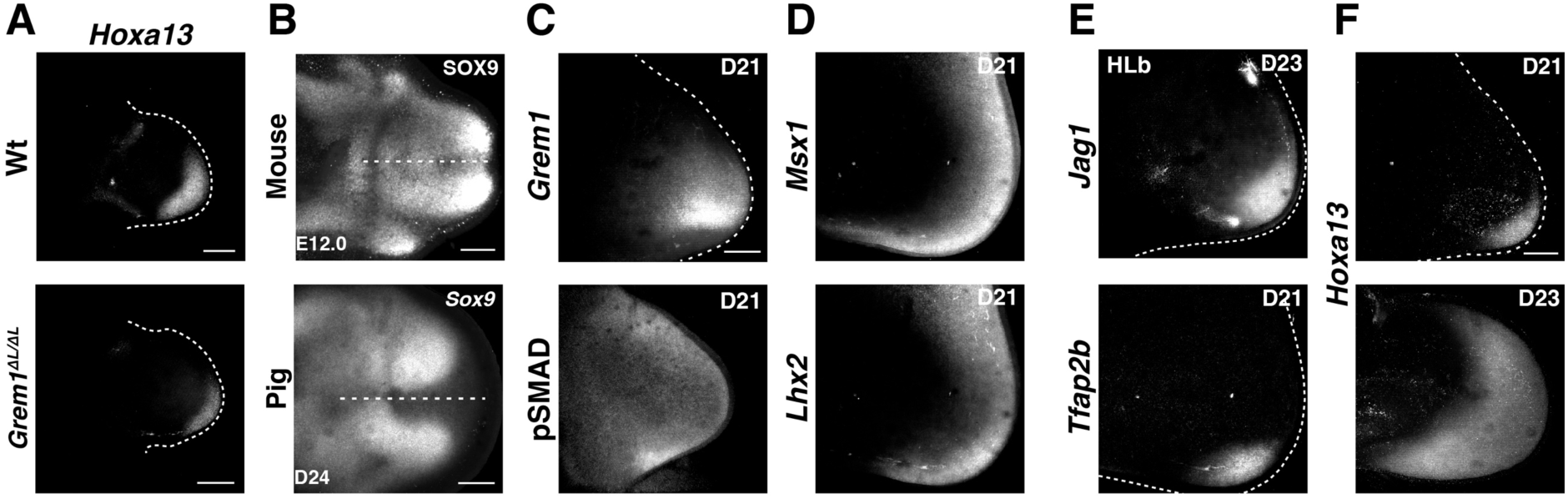
*Hoxa13* expression in *Grem1* and analysis of LMP signature genes in developmentally younger pig limb buds. (A) Hoxa13 expression in wild-type and Grem1 forelimb buds at E10.75. (B) Left panel: SOX9 protein distribution in *E1C5*^Δ/Δ^ tetradactyl mouse forelimb buds at E12.5 (n=6). Right panel: *Sox9* expression in pig forelimb buds at D24 (n=3). (C) Spatial distribution of *Grem1* (RNA-FISH) and pSMAD activity (immunofluorescence) in pig forelimb buds at gestational D21. (D) *Msx1* and *Lhx2* expression in pig forelimb buds at D21. (E) *Jag1* (hindlimb bud D23) and *Tfap2b* expression (forelimb bud D21) in pig limb buds. Note: pig hindlimb buds at D23 are equivalent to forelimb buds at D21(REF Tissiere). (F) The spatial distribution of *Hoxa13* in pig forelimb buds at D21 and D23. All analysis shown in C-F: minimally n=3 independent biological replicates analyzed. Scale bar: (A) 200µm, (B-F) 300 µm. All limb buds are oriented with anterior to the top and posterior to the bottom.

**in E1C5^Δ/Δ^ forelimb bud at E12.5.** Light sheet microscopy of an *E1C5*^Δ/Δ^ mouse

**E1C5 ^Δ/Δ^ forelimb bud at E12.5** Light sheet microscopy of an *E1C5*^Δ/Δ^ mouse forelimb

